# Intronised fluorophores improve reporter output and biosensor expression in plants

**DOI:** 10.64898/2026.09.21.753201

**Authors:** Sebastian Samwald, Tom Schreiber, Tonni Grube Andersen

## Abstract

Genetically encoded biosensors are central to plant cell biology, but low reporter signal can limit their use. Here, we developed modular intronised fluorophores to improve reporter output in transient and stable plant expression systems. Matched nuclear-localised triple mCitrine and mScarlet3 reporters were compared with and without introduced introns in *Nicotiana benthamiana* and *Arabidopsis thaliana*. Intronisation increased fluorescence, with the magnitude of enhancement depending on the fluorophore and expression context. Immunoblot analyses supported increased reporter accumulation. In Arabidopsis, intronised reporters driven by the tissue-specific *SCARECROW* promoter, produced stronger fluorescence signals that remained detectable farther from the root tip than non-intronised versions. Intronised versions of the abscisic acid (ABA) biosensor ABACUS2 retained ABA responsiveness and showed a lower frequency of reduced fluorescence between generations. These modular components provide a practical toolbox for improving fluorescent reporter detection and maintaining usable biosensor expression, supporting their application in plant cell biology and quantitative imaging.

## Introduction

Fluorescent proteins make it possible to visualise gene expression, protein localisation and physiological changes in living plant cells. For transcriptional reporters, weak or transient promoter activity can leave fluorescence close to background signals or autofluorescence. For genetically encoded biosensors, sufficient reporter abundance must be maintained across experiments and plant generations. The consequences of unstable expression are particularly apparent in fluorescence-based sensors, where multiple fluorescent proteins with similar sequences can lead to problematic expression or silencing. Examples of this include glucose nanosensors deployed in RNA-silencing mutants to overcome loss of expression, and successive generations of ABA sensors that have required careful evaluation of their behaviour in plants^1, 2^.

Reporter output can be increased by improving fluorophore properties, expressing tandem fluorescent proteins or concentrating the reporter in a defined subcellular compartment. Such setups have proved useful for resolving cell-type-specific patterns in plant development and signalling^3^. These strategies increase detectable fluorescence, but stable transgene expression remains sensitive to the properties of the expression construct and the transformed line^4^. A complementary approach is therefore needed to optimise the architecture of the reporter gene itself.

Introns can enhance transgene expression in plants, a phenomenon termed intron-mediated enhancement (IME)^5, 6^. Genome editing efficiency can be enhanced by intronisation of the Cas9 gene^7^. Intron splicing can also protect transgenes from RNA silencing. In *Arabidopsis thaliana* (thereafter Arabidopsis), introducing efficiently spliced introns into GFP reporter constructs reduced silencing and the production of RDR6-dependent secondary small interfering RNAs^8^. These findings provide a rationale for incorporating introns into fluorescent reporters, although the contribution of expression enhancement and protection from silencing must be evaluated in each construct and experimental context.

Established intronisation protocols and modular cloning systems provide a route to making this strategy accessible^9, 10^. Modular intron-insertion approaches have also been developed for *Chlamydomonas reinhardtii*^11^. Here, we apply intronisation to reusable fluorescent modules and test their performance in plant reporter applications. We first compare matched reporters in the rapid *N. benthamiana* transient expression system and followed by stable Arabidopsis transformants. We then examine a tissue-specific promoter and extend the approach to two ABACUS2 biosensor variants. This experimental progression tests whether intronised fluorophores improve reporter output in distinct expression contexts and help maintain useful biosensor expression across generations.

## Results

### Modular intronised fluorophores increase reporter output in transient expression assays

To initiate our analysis, we generated nuclear-localised triple mCitrine- and mScarlet3-based reporters with and without introduced introns. Following an established intronisation workflow, two plant introns were introduced into each fluorescent protein coding unit of the tandem reporters^10^. The corresponding intron-containing and intronless expression constructs otherwise shared the same design. Both reporter pairs were expressed under the Arabidopsis *ACT2* promoter and included an SV40 nuclear localization signal. This design allowed the effect of Intronisation to be assessed within each fluorophore pair. To compare reporter output in a transient expression system, we expressed the constructs in *N. benthamiana* leaves. A co-expressed nuclear-localised blue fluorescent reporter (tagBFP) provided an internal reference for fluorescence normalisation in the transient expression system. Both intronised and intronless constructs produced nuclear fluorescence in leaf epidermal cells (Fig. 1A). Normalized nuclear fluorescence was higher for the intronised versions of both reporters, with a more pronounced separation between the 3×mCitrine constructs than between the 3×mScarlet3 constructs (Fig. 1B). Immunoblot analyses provided an independent comparison of reporter accumulation. The HA-tagged fluorescent reporters and FLAG-tagged tagBFP were detected in extracts from the transiently expressing leaves (Fig. 1C). The immunoblot comparisons supported increased protein accumulation of the intronised reporters. Together, these results indicate that intronisation can increase reporter output in a leaf transient expression system.

**Figure 1:**
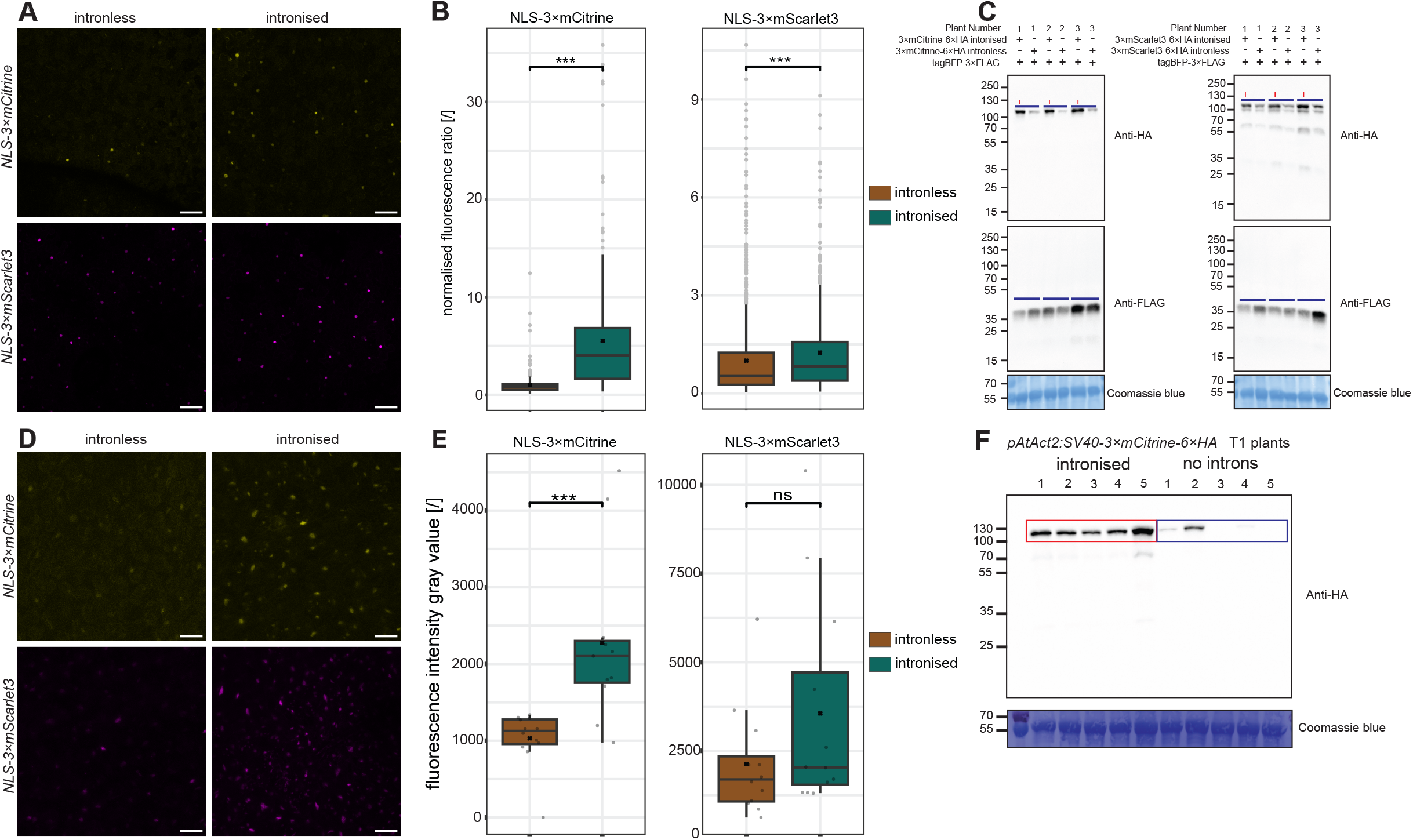
Intronised nuclear reporters increase fluorescence in transiently expressing *Nicotiana benthamiana* leavesand and in stable Arabidopsis transformants with fluorophore-dependent variation. (A) Representative nuclear fluorescence images of *pAtACT2:NLS-3×fluorophore-6×HA* of mCitrine and mScarlet3. Scale bars, 100 *µ*m (B) Nuclear mCitrine and mScarlet3 fluorescence normalized to the co-expressed blue fluorescent reference off the same expression plasmid. Number of images of independent leaves analysed is ≥10, resulting in number of nuclei of ≥229. Box plots: the line within the box marks the median, the box signifies the upper and lower quartiles, the whiskers represent the minimum and maximum within 1.5 × interquartile range, the × represents the mean. (C) Immunoblots detecting HA-tagged reporters and the FLAG-tagged tagBFP reference in triplicates. Each direct comparison of intronless and intronised constructs from the same plant has been indicated by a blue line. Coomassie staining is shown as a loading control. (D) Representative fluorescence images from independent T1 plants expressing nuclear-localised 3×mCitrine or 3×mScarlet3 under the ACT2 promoter. Intronised and intronless versions are shown for each reporter. Scale bars, 50 *µ*m. (E) Fluorescence intensity of nuclei comparisons across T1 transformants. Box plots: the line within the box marks the median, the box signifies the upper and lower quartiles, the whiskers represent the minimum and maximum within 1.5 × interquartile range, the × represents the mean. Number of images of independent T1 lines for each construct analysed is ≥9 (F) Anti-HA immunoblot of a randomly chosen subset of the displayed NLS-3×mCitrine lines (additionally tagged with 6×HA), representing five intronised and five intronless lines. Coomassie blue staining is shown as a loading control.

### Reporter enhancement varies between the tested constructs and expression contexts

We next tested *pACT2*-driven reporter pairs in independent T1 Arabidopsis transformants. Sampling independent insertion events allowed reporter performance to be evaluated across the variation encountered during routine transgenic line generation. Reporter fluorescence varied between lines in both the intronised and intronless groups (Fig. 1D, E). Intronised 3×mCitrine showed a clear increase in fluorescence, whereas the mScarlet3 comparison displayed greater between-line variation and a less pronounced separation between the intronless versus the intonised lines. For mCitrine-based constructs, the reporter accumulation was higher in the displayed intronised lines than in the intronless lines (Fig. 2C). Although intronised 3×mScarlet3-based constructs generated lines which showed an increased fluorescence above the intronless lines, this was not significantly different under the applied statististical constrains. Despite this, our results extend the reporter enhancement observed in transient assays to stable transformants, while showing that the magnitude and consistency of the effect vary between fluorophores, expression contexts, and each independent transformation event.

**Figure 2:**
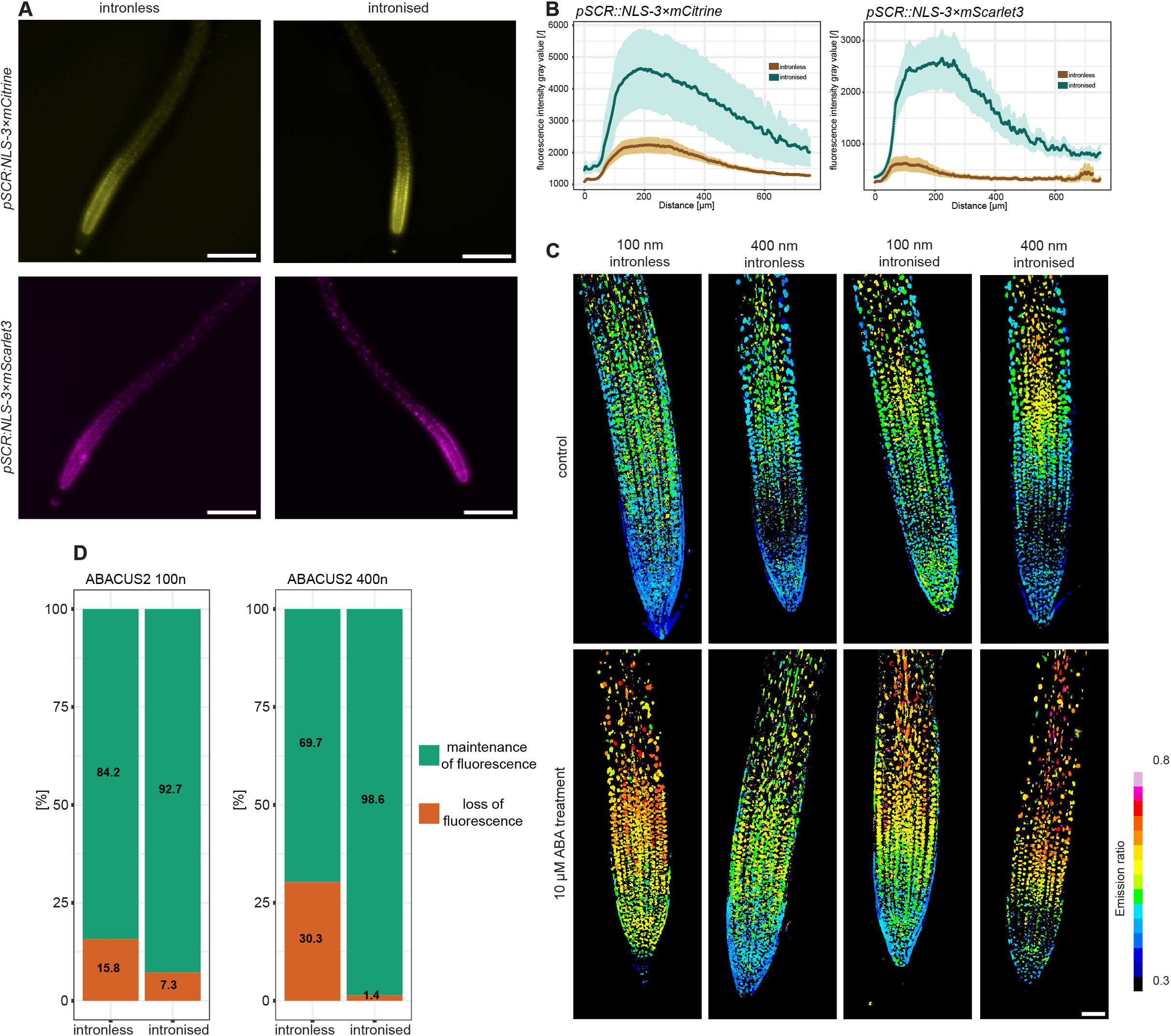
Intronised reporters produce stronger fluorescence under the SCR promoter and intronised ABACUS2 biosensors retain ABA responsiveness and show a lower frequency of fluorescence-defined expression loss. (A) Representative T1 roots expressing nuclear-localised intronised or intronless nuclear-localised 3×mCitrine and 3×mScarlet3 reporters under the SCR promoter. Scale bars, 250 *µ*m (B) Fluorescence profiles along the root axis for the same mCitrine (upper plot) and mScarlet3 (lower plot) fluorescence. The darker coloured lines indicate the mean, and the shaded regions indicate the SE. Number of images of independent T1 lines for each construct analysed is ≥13. (C) Representative root emission-ratio images for the intronless and intronised 100n and 400n ABACUS2 sensor variants under control conditions and after 24 h of treatment with 10 *µ*M ABA. The colour scale indicates the displayed emission ratio, with each line shifting its emission ratio when ABA treated; scale bar, 50 *µ*m. (D) Fractions assigned to the loss of fluorescence and maintenance of fluorescence categories in the source fluorescence screen of T3 plants. Any homozygous T3 line’s fluorescence intensity below 0.5× of the mean of its corresponding T2 parent line was counted has having lost its fluorescence. The fractions classified as having undergone loss of fluorescence were 15.8% and 7.3% for the intronless and intronised 100n variants, and 30.3% and 1.4% for the corresponding 400n variants, respectively. Number of analysed homozygous T3 roots of each construct analysed is ≥69.

### Intronised reporters increase fluorescence under a tissue-specific promoter

To assess if our reporter system affects developmental contexts where promoter activity reflects expression domains, we placed the nuclear-localised reporter pairs under the *SCARECROW (SCR)* promoter. *SCR* has an established role in root ground-tissue patterning, providing a well-characterized context for examining reporter output in the endodermal lineage^12^. We compared intronised and intronless reporters across independent transformation events and measured fluorescence along the root axis. For both 3×mCitrine and 3×mScarlet3, intronised reporters produced higher fluorescence profiles and retained detectable fluorescence farther from the root tip than the corresponding intronless reporters (Fig. 2A, B). These observations demonstrate enhanced reporter detection under the tested tissue-specific promoter, extending the region in which fluorescence could be measured under the imaging conditions used.

### Intronised ABACUS2 biosensors retain ABA responsiveness and show improved expression retention

As fluorescence-based biosensors are prone to silencing, we next tested whether intronised fluorophores could improve the performance and reliability. We introduced introns into the fluorescent protein coding regions of the two ABA sensitivity biosensor variants ABACUS2-100n and ABACUS2-400n, generating paired intronised and intronless constructs ^1^. During T1 screening, some strongly fluorescent seedlings showed severe growth impairment. Subsequent experiments used lines with detectable sensor expression and no obvious growth defects under the screening conditions. To assess ABA responsiveness, we compared reporter output after 24 h of treatment with 10 µM ABA and under control conditions. Both the intronised and intronless sensor variants showed ABA-responsive changes in the displayed ratio images (Fig. 2C). This comparison supports retention of ABA responsiveness following intronisation. We next compared fluorescence in T2 plants with that of their homozygous T3 progeny. The fraction of plants classified as loss of fluorescence (for this experiment defined as a drop in nuclear fluorescence signal intensity in T3 roots to below 0.5× of the mean of their segregating parental T2 line’s nuclear fluorescence signal intensity) in the fluorescence screen was lower for the intronised version of each sensor. For the 100n variant, the reported fractions were 15.8% without introns and 7.3% with introns; for the 400n variant, they were 30.3% and 1.4%, respectively (Fig. 2D). The fluorescence distributions for individual families provide additional information on the variation underlying these pooled comparisons (Fig. 2D, Fig. S2). These observations are consistent with improved retention of detectable biosensor expression between the T2 and T3 generations, particularly for the 400n variant.

## Discussion

This study evaluates intronisation as a practical means of improving fluorescent reporter output in plants. Matched reporter comparisons in transient *N. benthamiana* assays, independent Arabidopsis transformants and a tissue-specific expression context show that intronised fluorescent modules can increase the signal available for imaging. The ABACUS2 comparisons further suggest that the approach can help maintain detectable biosensor expression across generations. Packaging these fluorophores as reusable modules allows their benefits to be tested within established cloning workflows, thereby reducing the need to redesign the fluorescent coding sequence for each application.

The magnitude of reporter enhancement depends on context. The two fluorophores did not show equally strong separation in every assay, and independent stable lines retained substantial variation. Intronisation should consequently be evaluated as one component of construct design rather than assumed to confer a fixed increase in output. The present comparisons also do not establish whether greater reporter accumulation results primarily from changes in transcript abundance, RNA processing, translation or protection from RNA silencing. Previous work demonstrated that efficient intron splicing can reduce RDR6-dependent silencing in Arabidopsis^8^, providing a plausible explanation for part of the effect without establishing that it accounts for the differences observed here.

For transcriptional reporters, increased fluorescence signal improves the ability to detect accumulated reporter activity, but its biological interpretation remains dependent on the expression system. The *SCR* profiles illustrate this distinction as intronised reporters were brighter and remained detectable farther along the root. It is important to mention that these observations could reflect higher reporter accumulation, continued low-level expression or changes in expression associated with the introduced introns. They do not establish an extended endogenous transcriptional domain. Although onset of expression is not changed, a higher protein abundance could possibly lead to longer persistence fluorescence. Reporter maturation and persistence must therefore be considered when using the constructs to infer current promoter activity. Similarly, an increase in fluorescence intensity does not specify the improvement in signal-to-noise ratio without measurements of the relevant background and noise.

Overall, intron insertion enhanced 3×mCitrine and 3×mScarlet3 reporter output to different extents, as observed in transient expression in *N. benthamiana* (Fig. 1A, B) and stable expression in *Arabidopsis* (Fig. 1D, E, F). This variation could reflect differences in intron identity, position and coding-sequence context. The 3×mScarlet3 construct includes three introns newly incorporated into our reporter designs (AtPGK-i3, AtPGK-i5 and AtTOPII-i1). Genomic analyses show that the median intron length is shorter in *Arabidopsis* than in maize and rice^13^, while a minimum functional length of 70–73 nucleotides (nt) was identified for a synthetic intron tested in monocot and dicot protoplasts^14^. These findings provide context for considering intron length during construct design. However, the lengths of AtPGK-i5 (77 nt) and AtTOPII-i1 (175 nt) alone do not establish whether either intron limits reporter expression. As neither splicing efficiency nor the contributions of individual introns were assessed here, the basis of the limited enhancement of ACT2-driven 3×mScarlet3 remains unresolved. Newly generated constructs should therefore be evaluated in their intended expression context by comparing intronised modules with matched intronless counterparts. Initial transient expression can support this evaluation, followed by testing and, where necessary, optimisation in the intended stable expression system.

The biosensor experiments highlight both the benefits and the constraints of increasing reporter abundance. The reduced frequency of fluorescence-defined expression loss is practically useful because maintaining sensor expression is necessary for reproducible imaging. However, brighter starting expression could also change the fraction of plants falling below the detection threshold. Insertion-line effects, transgene segregation, and the selection of viable expressing lines must therefore be considered when interpreting comparisons across generations. Direct evidence for altered silencing would be needed to assign the observed expression retention to a specific molecular pathway.

Increasing biosensor abundance can also perturb the process being measured. The growth defects observed in some strongly fluorescent T1 seedlings emphasize the need to select and characterize lines with an appropriate expression level. Their cause was not established in this study. ABA sequestration is one possible explanation, but ABACUS2-expressing plants have also been reported to show ABA hypersensitivity associated with residual activity of the sensor receptor component^1^. The affected seedlings therefore cannot be used to infer depletion of endogenous ABA. Likewise, the absence of an obvious growth defect under standard conditions does not demonstrate that hormone responses are unchanged.

Taken together, our findings support that intronised fluorophores can be incorporated into reporter constructs while other components, including biosensor recognition domains, could continue to be developed independently. By providing matched intronised and intronless modules, this resource enables those direct comparisons and offers a practical route to improving fluorescent reporter detection and the maintenance of usable biosensor expression in plants.

## Materials and methods

### Plant material, growth conditions and transformation

#### Transient expression in *N. benthamiana*

*N. benthamiana* plants were cultivated in a greenhouse cabin exposed to ambient sunlight and supplemented with a broad-spectrum 16 h photoperiod from LED lights. The pots are irrigated daily with a nutrient solution containing an electrical conductivity of 2.2 mS/cm; pH 5.6; potassium (0.46 mmol/L); calcium (0.38 mmol/L); magnesium (0.16 mmol/L); nitrogen-to-potassium ratio of 1.8; ammonium fraction of total nitrogen of 0.05 and phosphorus ratio of 0.055. *Agrobacterium tumefaciens GV3101* (carrying the helper plasmid pMP90)^15^, carrying the desired construct was cultured overnight and twice washed and resuspended to a final OD_600_ of 0.3 in 0.01 M 2-(N-morpholino)ethanesulfonic acid (MES) pH 5.6, 0.01 M MgCl2, 0.01 µM acetosyringone. No P19 silencing suppressor was added. Intronised versus intronless constructs were syringe-infiltrated into respective leaves of the same individual 4-week-old *N. benthamiana* plants. Material for experiments was harvested two days post-infiltration, as 6 leaf discs per sample using a cork borer n°4 (diameter 8.8 mm).

#### Arabidopsis seed sterilisation and sterile growth

Arabidopsis seeds were placed in an exicator, together with a glass beaker containing 100 mL Sodium hypochlorite (12% Cl, stabilised, Roth). Chlorine gas was produced by addition of 3 mL HCl (37%) before immediately closing the exicator. The seeds were exposed to the chlorine gas for 3 h.

For all seedling assays and sterile plant growth, seeds were placed on ½ strength Murashige & Skoog medium incl. MES buffer (Duchefa Biochemie, M0254.0010) media at pH 5.8 with 0.8% (w/v) plant agar (Duchefa Biochemie, P1001.1000) in square plates (12×12cm). After two days of stratification in the dark at 4°C, the plates were moved into an upright but slightly angled position to grow at 21°C under a 16-h-light/8-h-dark photoperiod.

### Arabidopsis transformation

All Arabidopsis lines used in this work are in the Col-0 background. Stable genetic transformation of Arabidopsis has been carried out using the floral dip method^16^. In brief: A single colony of *Agrobacterium tumefaciens* (GV3101) transformed with the desired plasmid (Supplementary Table1) was inoculated into 10 mL LB liquid culture and grown overnight with the additional appropriate selective antibiotics. 1-2 mL of this culture was used to inoculate 200 mL LB, including the appropriate antibiotics. After a further day of growth, the cultures were centrifuged for 15 min., at 5422 × g, discarded the supernatant and resuspended the pellet in 200 mL infiltration media (2.165 g/L Murashige and Skoog (MS) medium basal salt mix, 50 g/L Sucrose), by shaking. Silwet L-77 (50 μL per 200 mL) of was added to the bacterial suspension mixed manually. Nine Arabidopsis plants grown in a 9 cm diameter pot, were dipped for 45-60 s in the bacterial solution, and kept in the dark overnight.

### Arabidopsis transgenic seed selection

Transgenic T1 Arabidopsis seeds were identified by FastRed seed coat fluorescence. Independent T1 transformants were used for all reporter comparisons. FastRed fluorescence of T2 seeds was used to assess their mendelian ratio, to avoid insertion lines with multiple insertion sites. T3 seeds were similarly assessed for homozygosity by FastRed fluorescence.

### Transient expression and fluorescence normalisation

Reporter constructs were transiently expressed in *N. benthamiana* leaf epidermal cells. The transient expression plasmids also carried a cassette encoding a nuclear-localised blue fluorescent reference. This reference was used to normalize reporter fluorescence within individual cells.

### Normalisation of fluorescence ratios

In transient expression assays of *N. benthamiana* each plant cell can exhibit a different strength of reporter expression due to its independent transformation. To account for this, we have used reporter plasmids which not only carry *pAtACT2:NLS-3×fluorophore-6×HA* (mCitrine or mScarle3) but additionally within the same plasmid another constitutively expressed fluorophore: *pAtUBQ10::NLS-tagBFP-3×FLAG*. Using FIJI^17^ based on the blue fluorescence channel, we used the circle tool to manually select all nuclei, and export their mean grey value for both the tagBFP and mCitrine or mScarlet3 fluorescence respectively. For each individual nucleus we determined the fluorescence ratio by dividing the mCitrine or mScarlet3 channel mean grey value by the tagBFP mean grey value. Each datapoint was divided by the mean value of the intronless control to further normalise the ratio.

### Fluorescence of Arabidopsis nuclei

Nuclear fluorescence of roots of independent T1 Arabidopsis transformation lines was detected from a z-projection using a FIJI^17^ macro detecting objects with a minimum size of 60 pixels, a maximum size of 2000 pixels, a circularity shape factor of 0.1, after a thresholding based on the ‘Triangle’ method, Gaussian blur application with a sigma of 1, and watershedding. The mean of all nuclei imaged and detected from one root of one independent transformation line, was used as a single data point.

### Construct design and modular cloning

Constructs for Arabidopsis and *N. benthamiana* transformation and expression were generated using the golden gate cloning system^9, 18^. Two plant introns were inserted into each fluorescent protein coding unit following the intronisation strategy described by Schreiber and Marillonnet ^10^, using introns selected on the basis of previous plant expression work^6^. Paired triple mCitrine and mScarlet3 reporters were codon optimised for Arabidopsis (VectorBuilder, excluding BsaI, BpiI, and EspIII sites) designed with and without introduced introns, for different golden gate cassettes, as well as versions with and without the nuclear localisation signal SV40 (exon length between 300 and 400nt). The majority of introduced introns have been shown to be effective in increasing SpCas9 expression levels in Arabidopsis^7^. In addition, four new introns were incorporated: one hybrid intron in mCitrine (AtCSLA07 (At2g35650) Intron 4 (5’, 40nt) fused to AtMOR1 (At2g35630) Intron 25 (3’, 69nt)) and three natural introns in mScarlet3 (AtPGK (At3g12780) Intron 3 (96nt) and intron 5 (77nt) and AtTOPII (At3g23890) Intron 1 (175nt)) (Fig. S1). Similarly, two introns were integrated into edCitrineT9 and the T7edCerulean fluorophore components of ABACUS2 respectively (Fig. S1). All splice sites were optimised to achieve a splice confidence of 1.00 or as close as possible thereof, and any potential cryptic splice sites were adapted with silent mutations, using the using the NetGene2 server soft-ware (https://services.healthtech.dtu.dk/services/NetGene2-2.42/^19^)^20^. Complete and parts of the intronised fluorophore constructs were generated using gene synthesis (Genscript). Parts were assembled into plasmids following golden gate assembly logic (Supplementary Table 4, 5, and 8).

Coding sequences were generated to be either flanked by the pAtACT2, the pSCR, or the pUBQ10 promoters and their corresponding native terminators. Each expression plasmid carried the OLESIN coding sequence tagged with TagRFP flanked by the Arabidopsis native OLESIN promoter and terminator to allow for FastRed seed selection^21^.

Modular constructs followed the plant MoClo framework^9^. Cloning strategies, all used golden gate parts, and final plasmids are listed in the supplementary tables 1-8. Cloning PCR were carried out following the Bounce PCR workflow ^22^ using the PrimeSTAR® GXL Premix (Takara, R051A). PCR products were size assessed by gel electrophoresis, and the correctly sized DNA bands were extracted using QIAquick Gel Extraction workflow (Quiagen, 28704). Plasmid preps were carried out using the QIAprep Spin Miniprep workflow (Quiagen, 27104). For both gel extractions and plasmid preps EconoSpin® All-In-One DNA Only Mini Spin Column (Epoch Life Science, 1920-250) were used.

### Microscopy and root fluorescence profiles

Nuclear reporter fluorescence was imaged in transiently expressing leaves and stable *Arabidopsis* transformants. Confocal imaging of Arabidopsis and *N. benthamiana* performed using a Zeiss LSM 980 confocal microscope. tagBFP was excited at 405 nm, and emission was detected at 423–482 nm; mCitrine was excited at 488 nm, and emission was detected at 525-587 nm. mScarlet3 was excited at 561 nm, and emission was detected at 587-633 nm.

ABACUS2 fluorescence was detected by sequential scanning with a 445 nm laser was used to excite the edCerulean (for edCerulean and edCitrine FRET emission) and 514 nm lasers were used to excite edCitrine (for edCitrine emission, acting as an expression control). Emission settings were 460–500 nm for Cerulean and 525–560 nm for edCitrine.

For the *SCR*-driven reporters, fluorescence was quantified as a function of position along the root axis using the segmented line tool with a line width of 100 µm placed centrally along the root in FIJI^17^ starting from the root tip in a maximum z-projection. Arabidopsis roots were imaged using a Zeiss Axio Zoom.V16 stereomicroscope equipped with a Zeiss Axiocam 705 mono or colour camera.

### Protein extraction and immunoblotting

To prepare for protein extraction, either 6 leaf discs of *N. benthamiana* or 2 mature leaves of Arabidopsis were harvested per genotype and per replicate in a 1.7 mL centrifugation tube together with two metal beads (∅ 3.2 mm) and frozen in liquid nitrogen. The samples were ground using a bead mill (Qiagen, TissueLyserII) at 30 Hz for 1 min, and extracted in 500 μL adapted Laemmli buffer (0.13 M Tris-HCl pH 6.8, 4% SDS (Sodium Dodecyl Sulfate), 20% Glycerol, 0.2 M DTT (Dithiothreitol) and 0.02% Bromophenol blue). After 30 min continuous tumbling at 4°C, the samples were centrifuged at 4°C at 18213 rcf for 15 min. The supernatant’s protein concentration was assessed using Pierce™ 660 nm Protein-Assay-Reagent (Thermo Fisher Scientific 22660) together Ionic Detergent Compatibility Reagent for Pierce™ 660nm Protein Assay Reagent (Thermo Fisher Scientific 22663) to equalise loading. The supernatant was heated for 15 min at 95°C, and volume calculated for equal loading was loaded on 10% polyacrylamide gels (BIORAD, 10% Mini-PROTEAN® TGX™ Precast Protein Gels, 15-well). After wet transfer (BIORAD, Mini Trans-Blot system), the membranes (BIORAD, Immun-Blot PVDF Membrane) were blocked for 1h at room temperature, and probed overnight at 4°C with the corresponding antibodies, α-FLAG-HRP (abcam, ab 49763, 1 in 5000) or α-HA-HRP (abcam, ab173826, 1 in 5000), after four 5 min washes in TBST, we detected for chemiluminescence signals in a ChemiDoc MP Imaging System (BIORAD) using enhanced chemiluminescent (ECL) substrate (Thermo Scientific, SuperSignal™ West Femto Maximum Sensitivity Substrate).

Additional Coomassie Brilliant Blue (CBB) staining to confirm equal loading ^23^ was carried out on the same membranes, by briefly floating each membrane in CBB staining buffer [250 mL dH_2_O, 200 mL Methanol, 50 mL Acetic Acid, 1.25 g CBB powder (SERVA Blue G, COOMASSIE® Brilliant Blue G-250)], and destained over multiple hours under gentle agitation in destaining buffer [500 mL dH_2_O, 400 mL Methanol, 50 mL Acetic Acid], before a final wash in dH_2_O and drying before imaging.

### ABACUS2 line selection and ABA treatment

During T1 screening, some seedlings with strong ABACUS2 fluorescence showed severe growth impairment, and never continued growth past a seedling stage nor managed to develop true leaves. Subsequent experiments used lines without obvious growth defects under the screening conditions. To assess ABA responsiveness seven-day old seedlings were transferred onto fresh plates of ½ MS media containing either 10 µM ABA (Duchefa Biochemie, A0941) or the corresponding Ethanol treatment control. After 24 h continued growth the seedlings were prepared on microscopy slides and the root tip regions of each seedling promptly imaged as described below. Images were analysed for their nuclear emission ratios using FRETENATOR2^24 25^.

### Expression retention between generations

Two independent genetic transformation lines of each construct of both intronised and intronless versions of the 100n and 400n ABACUS2 biosensor were selected to continue downstream analysis. From T2 seeds, plants which germinated whose seed coats exhibited FastRed signals were selected to generate T3 seeds. In total T3 seeds from 10 different T2 plants of each line were collected. These T3 seeds were sown out together with seeds of their parental T2 seeds. Roots of these plants were imaged in one session with the exact same microscopy settings. edCitrine was excited with a 514 nm laser and its emission detected at 525–560 nm. Each nuclei’s fluorescence was detected from a z-projection using a FIJI ^17^ macro detecting objects with a minimum size of 20 pixels, a maximum size of 250 pixels, a circularity shape factor of 0.2, after a double thresholding based on the ‘Otsu white stack’ followed by the ‘Shanbhag white stack’ method. The mean of all nuclei imaged and detected from one root, was used as a single data point. All T3 and T2 data points were normalised to the mean nuclei fluorescence of all T2 roots imaged from their corresponding independent transformation line.

Homozygous T3 seeds were determined by FastRed signal in all seeds of the same T2 plant origin. These data points were directly compared to the mean fluorescence of their parental T2. If they exhibited a value below 0.5× of the mean of their parental T2 plant, they were counted as having lost their fluorescence signal, whilst if they exhibited a value above 0.5× of the mean of their parental T2 plant they were counted as having retained their fluorescence.

## Statistical analysis

Statistical analyses were performed using R 4.6.1 (Happy Hop).

Normalised nuclear fluorescence intensity values of datasets with between only two groups we used the non-parametric Wilcoxon. For comparing the T2 ABACUS2 as the control group with each T3 ABACUS2 line’s fluorescence we used the non-parametric post-hoc test Dunn’s test. p values were corrected for multiple comparisons using the Holm-Bonferroni method. ns indicates non-significance, p.adj < 0.001 ∼ ***, p.adj < 0.01 ∼**, p.adj < 0.05 ∼ *.

## Data and materials availability

The intronised and corresponding intronless fluorophore modules will be deposited for distribution on Addgene upon publication. Correspondence and requests for materials should be addressed to Tonni Grube Andersen. The planned collection includes nuclear-localized complete coding-sequence modules and N- and C-terminal fusion modules compatible with the plant MoClo system.

## Author contributions

Conceptualization: S.S., T.S., T.G.A.; Funding acquisition: S.S., T.G.A.; Investigation: S.S.,T.S.; Methodology: S.S., T.S., T.G.A.; Project administration: S.S., T.G.A.; Supervision: T.G.A.; Visualization: S.S., T.G.A.; Writing – original draft: S.S., T.G.A.; Writing – review & editing: all authors.

## Acknowledgements and funding

We thank Aristeidis Stamatakis and the greenhouse team at MPIPZ for help with plant growth, and Anika Schröder for the help with plant husbandry and genetic transformation. We thank James Rowe for providing plasmids and insightful discussions. We thank all the members of the TGA group in Cologne and Zürich for their assistance with this project.

This study received funding from the Sofja Kovalevskaja program at the Alexander von Humboldt foundation (T.G.A.); the Max Planck Society (T.G.A.); the Deutsche Forschungsgemeinschaft (DFG, German Research Foundation) under Germany’s Excellence Strategy – EXC-2048/1 – project ID 390686111 (S.S. and T.G.A.); the European Union’s Horizon 2020 research and innovation programme under the Marie Skłodowska-Curie grant agreement 101152197 (S.S.).

---

**Fig. S1: Graphical representation of fluorophore intronisation**

(A) 3×mCitrine intronised, (B) 3×mScarlet3 intronised, (C) edCitrineT9, (D) T7edCerulean. The colours indicate as follows: yellow: exons, grey: introns, red: branch points, blue: Donor splice site, light yellow triangles: Acceptor splice sites, pink: domestication for golden gate, to achieve efficient splicing, removal of potential cryptic splice site.

**Fig. S2: Fluorescence distributions for individual ABACUS2 lines comparing T2 with their direct descendant T3 lines.**

Two independent transformation events per intronless and intronised construct of ABACUS2 sensor variants 100n and 400n have been taken forward into their T3 generation. Root nuclear fluorescence of the T2 line and their corresponding T3 progeny have were compared directly and normalised to their insertion specific T2 fluorescence intensity mean. One data point consists of the mean nuclear fluorescence intensity of one root. All data points below 0.5× of their insertion specific T2 fluorescence intensity mean are indicated in red, whilst all above this threshold are indicated in black. Any data point below this threshold was counted as having undergone loss of fluorescence, whilst all above were counted as having maintained their fluorescence from the T2 generation to the T3 generation. Number of individual roots per box plot is ≥10.

Orange indicates T2 segregating parental line, magenta indicated heterogzygous T3 lines and green indicates homozygous T3 lines according to their FastRed seedcoat ratios. A Dunn test was carried out to compare all T3 lines to the T2 parental line as the reference group with a Bonferroni correction. Any statistical significance was indicated (adjusted p-values: < 0.001 ∼ ***, < 0.01 ∼ **,< 0.05 ∼ *), whilst all non-significant comparisons were not indicated.

