## Supplementary Figures, Extended Data, Supplementary Tables for "Intronised fluorophores improve reporter output and biosensor expression in plants"

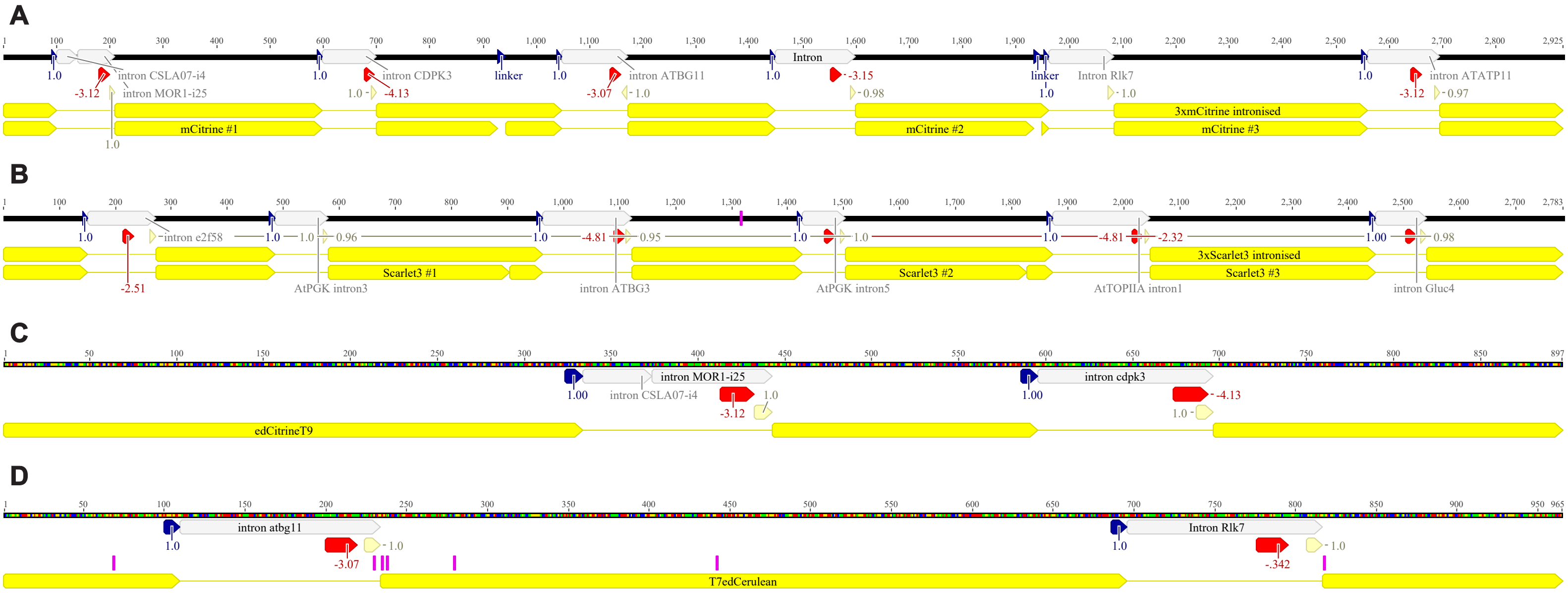


**Fig. S1: Graphical representation of fluorophore intronisation**

(A) 3×mCitrine intronised, (B) 3×mScarlet3 intronised, (C) edCitrineT9, (D) T7edCerulean. The colours indicate as follows: yellow: exons, grey: introns, red: branch points, blue: Donor splice site, light yellow triangles: Acceptor splice sites, pink: domestication for golden gate, to achieve efficient splicing, removal of potential cryptic splice site.

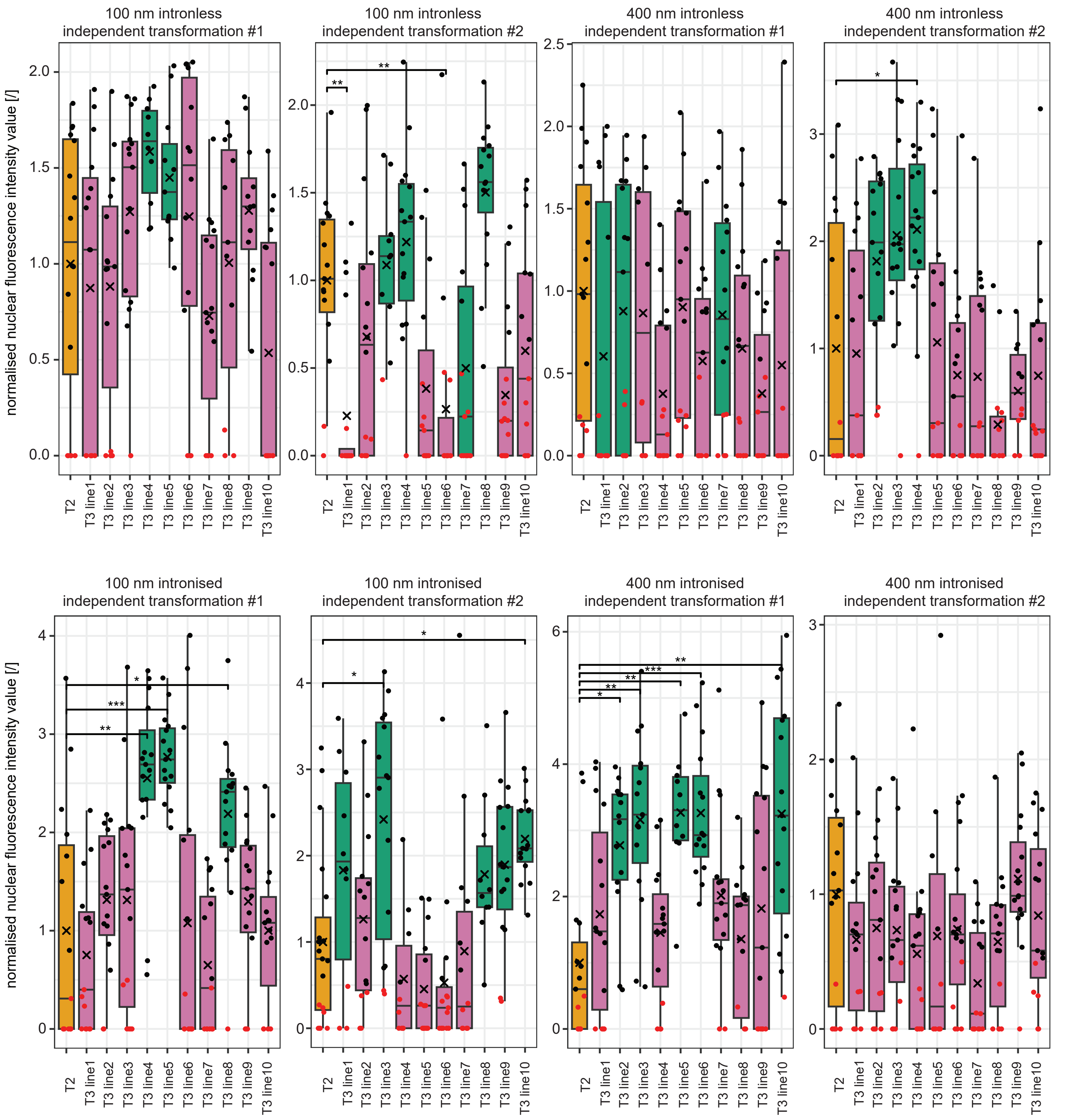


**Fig. S2: Fluorescence distributions for individual ABACUS2 lines comparing T2 with their direct descendant T3 lines.**

Two independent transformation events per intronless and intronised construct of ABACUS2 sensor variants 100n and 400n have been taken forward into their T3 generation. Root nuclear fluorescence of the T2 line and their corresponding T3 progeny have were compared directly and normalised to their insertion specific T2 fluorescence intensity mean. One data point consists of the mean nuclear fluorescence intensity of one root. All data points below 0.5× of their insertion specific T2 fluorescence intensity mean are indicated in red, whilst all above this threshold are indicated in black. Any data point below this threshold was counted as having undergone loss of fluorescence, whilst all above were counted as having maintained their fluorescence from the T2 generation to the T3 generation. Number of individual roots per box plot is ≥10. Orange indicates T2 segregating parental line, magenta indicated heterogzygous T3 lines and green indicates homozygous T3 lines according to their FastRed seedcoat ratios. A Dunn test was carried out to compare all T3 lines to the T2 parental line as the reference group with a Bonferroni correction. Any statistical significance was indicated (adjusted p-values: $< 0.001$ ~ ***, $< 0.01$ ~ **,$< 0.05$ ~ *), whilst all non-significant comparisons were not indicated.

3×mCitrine (exon, *intron*)

ATGGTGAGCAAGGGCGAAGAGTTGTTCACTGGTGTTGTTCCTATCCTCGTTGAGCTTGACGGTGATGT
GAACGGGCATAAGTTCTCCGTTTCTGGTGAAGGTAACATTCCTTAGTTACCTTTCTTTTCTTTTTCCA
TCATCTATCAATTTCTTTGCGGAAATTTATTTGAAGCTGTAGAGTTAAAATTGAGTCTTTTAAACTTT
TGTAGGTGAGGGAGATGCTACTTACGGAAAGCTCACCCTCAAGTTCATCTGTACCACTGGAAAGCTCC
CTGTGCCTTGGCCTACTCTCGTTACTACTTTCGGATACGGGCTCATGTGCTTCGCTAGATACCCTGAT
CATATGAAGCAGCACGACTTCTTCAAGAGCGCTATGCCTGAGGGATACGTGCAAGAGAGAACCATTTT
CTTCAAGGACGACGGGAACTACAAGACCAGAGCTGAGGTTAAGTTCGAAGGTGACACCCTCGTGAACA
GGATCGAGCTTAAGGGCATCGACTTCAAAGAGGACGGAAACATCCTCGGACACAAGCTCGAGTACAAC
TACAACAGCCACAACGTGTACATCATGGCCGACAAGCAGAAGAACGGCATCAAGGTAAGTTGTTACTT
ATGATTGTTTTCCTCTCTGCTACATGTATTTTGTTGTTCATTTCTGTAAGATATAAGAATTGAGTTTT
CCTCTGATGATATTATTAGGTCAACTTCAAGATCAGGCACAACATCGAGGACGGATCTGTTCAGCTCG
CTGATCATTACCAGCAGAACACCCCTATTGGAGATGGACCTGTTCTTCTCCCTGACAACCACTACCTC
AGCTACCAGTCTAAGCTCAGCAAGGACCCTAACGAGAAGAGGGATCATATGGTGCTCCTCGAGTTCGT
TACTGCTGCTGGAATCACTCTCGGAATGGACGAGCTTTACAAAGGCGGTGGTGGATCCATGGTGTCCA
AGGGTGAAGAACTTTTCACAGGTGTGGTGCCGATCCTTGTGGAACTCGATGGTGATGTCAATGGCCAC
AAGTTCAGCGTGAGCGGAGAAGGCGAAGGTAAATCCTGGTCCACACTTTTACGATAAAAACACAAGAT
TTTAAACTATGAACTGATCAATAATCATTCCTAAAAGACCACACTTTTGTTTTGTTTCTAAAGTAATT
TTTACTGTTATAACAGGTGATGCAACATATGGTAAGTTGACGCTGAAGTTTATCTGTACGACCGGGAA
GTTGCCTGTTCCATGGCCAACACTTGTGACCACATTCGGCTACGGATTGATGTGTTTCGCTCGTTACC
CAGACCACATGAAGCAACATGATTTCTTTAAGTCCGCCATGCCAGAGGGCTATGTCCAAGAAAGGACG
ATCTTTTTCAAGGATGATGGCAATTATAAGACCCGTGCCGAAGTGAAATTCGAGGGCGATACTCTTGT
GAACCGTATCGAGTTGAAAGGTAAGTTCTGCATTTGGTTATGCTCCTTGCATTTTAGGTGTTCGTCGC
ACTTCCATTTCCATGAATAGCTAAGATTTTTTTCTCTGCATTCATTCTTCTTGCCTCAGTTCTAACT
GTTTGTGGTATTTTTGTTTTAATTATTGCTACAGGTATTGATTTTAAAGAGGATGGAACATTTTGGG
GCACAAGTTGGAGTATAATTACAACTCGCATAACGTCTACATTATGGCGGATAAGCAAAAGAATGGTA
TCAAAGTGAATTTTAAGATCCGTCATAACATCGAAGATGGCAGCGTGCAGTTGGCAGATCACTACCAA
CAAAACACGCCGATCGGTGATGGTCCTGTGCTTCTTCCAGATAACCATTACCTGTCCTACCAGAGCAA
GCTCTCTAAGGATCCGAATGAGAAGCGTGATCACATGGTCCTTTTGGAGTTTGTGACCGCTGCTGGTA
TTACCCTTGGCATGGATGAACTCTACAAAGGCGGCGGAGGTAGCATGGTTTCTAAAGGTAAAGCAACT
GTGTTTTAATCAATTTCTTGTCAGGATATATGGATTATAACTTAATTTTTGAGAAATCTGTAGTATTT
GGCGTGAAATGAGTTTGCTTTTTGGTTTCTCCCGTGTTATAGGTGAGGAACTCTTTACCGGGGTCGTG
CCTATTCTTGTCGAGCTTGATGGGGACGTTAACGGCCACAAATTCTCGGTGTCAGGTGAAGGTGAAGG
CGACGCTACCTATGGGAAGCTCACTTTGAAATTCATATGTACTACGGGCAAGCTGCCGGTTCCTTGGC
CGACTTTGGTTACAACATTTGGCTATGGCCTGATGTGCTTTGCCAGGTATCCGGACCATATGAAGCAG
CATGATTTCTTCAAAAGTGCCATGCCGGAAGGTTATGTGCAAGAGCGTACGATATTCTTTAAAGATGA
CGGTAATTACAAGACGAGGGCCGAAGTCAAGTTTGAAGGGGATACACTGGTCAACCGAATCGAACTCA
AGGGAATAGATTTCAAAGAAGATGGTAATATCCTGGGGCATAAGCTAGAGTATAACTATAATTCCCAC
AATGTCTATATCATGGCTGACAAACAAAAGAACGGTATTAAGGTAAAGTTTCCAACTTTCCTTTACCA
TATCAAACTAAAGTTCGAAACTTTTTATTTGATCAACTTCAAGGCCACCCGATCTTTCTATTCCTGAT
TAATTTGTGATGAATCCATATTGACTTTTGATGGTTACGCAGGTGAACTTCAAAATCCGACATAATAT
TGAGGACGGCAGCGTCCAGCTTGCTGACCACTATCAACAGAATACCCCAATCGGTGACGGACCAGTTC
TCTTGCCTGATAATCACTACTTGTCGTACCAGTCCAAGTTGTCGAAAGATCCTAACGAAAAGCGTGAC
CACATGGTGTTGCTCGAGTTTGTCACAGCAGCAGGTATCACGCTCGGTATGGATGAGTTGTATAAGTG
A

3×mScarlet3 (exon, *intron*)

ATGGATTCTACCGAGGCCGTGATCAAAGAATTCATGAGGTTCAAGGTGCACATGGAAGGCAGCATGAA
CGGACATGAGTTCGAGATCGAAGGTGAAGGTGAGGGTAGACCTTACGAGGGAACTCAGACTGCTAAGC
TCAGAGTGACAAAAGGTCTGTCTTTCCTATTTCATATGTTTAATCCTAGGAATTTGATCAATTGATTG
TATGTATGTCGATCCCAAGACTTTCTTGTTCACTTATATCTTAACTCTCTCTTTGCTGTTTCTTGCAG
GTGGACCGCTTCCGTTCTCTTGGGATATCCTTTCACCGCAGTTCATGTACGGCTCTAGGGCTTTTACT
AAGCACCCTGCTGATATCCCGGACTACTGGAAGCAATCTTTCCCAGAGGGATTCAAGTGGGAGCGTGT
GATGAATTTCGAGGATGGTGGTGCTGTTTCTGTGGCTCAGGATACTTCTCTTGAGGACGGAACCCTCA
TCTACAAGGTAATATTTTGATAGAGATTATTCTATTCAGCATTGAAGCATCGTTGCTACAAATGGTAC
CATGAACTTTTTGTTTACTTCTCTCTACCTCTATAGGTTAAGCTCCGTGGAACTAACTTCCCTCCTGA
TGGACCTGTGATGCAGAAAAAGACCATGGGCTGGGAAGCTTCTACTGAGAGACTTTACCCTGAGGACG
TTGTGCTCAAGGGCGATATCAAGATGGCTCTCAGACTTAAGGACGGTGGACGTTACCTCGCTGATTTC
AAGACTACCTACAGGGCCAAGAAACCTGTGCAAATGCCTGGGGCTTTCAACATCGATAGGAAGCTCGA
TATCACCAGCCACAACGAGGATTACACTGTGGTTGAGCAGTACGAGAGATCTGTGGCTAGACACTCTA
CTGGTGGAAGCGGAGGATCTATGGATAGCACTGAGGCTGTCATCAAAGAATTCATGCGTTTCAAAGTT
CATATGGAAGGTAAAATATTGGATGCCAGACGATATTCTTTCTTTTGATTTGTAACTTTTTCCTGTCA
AGGTCGATAAATTTTATTTTTTTTGGTAAAAGGTCGATAATTTTTTTTTGGAGCCATTATGTAATTTT
CCTAATTAACTGAACCAAAATTATACAAACCAGGTTCCATGAATGGGCACGAATTTGAAATTGAAGGC
GAAGGCGAGGGACGTCCGTATGAAGGTACTCAAACAGCTAAGCTGAGGGTTACCAAAGGCGGTCCTCT
TCCTTTTTCCTGGGATATCTTGAGCCCTCAATTCATGTATGGCAGCAGGGCCTTCACCAAACATCCAG
CAGATATCCCTGATTATTGGAAACAAAGCTTCCCGGAAGGGTTTAAGTGGGAGAGAGTCATGAACTTC
GAAGATGGCGGAGCAGTGAGTGTTGCTCAAGACACTTCTTTGGAGGATGGGACTCTGATCTATAAGGT
AAGAAAATGATTCCTCATAGAGTAAACAAATGGTAGATGAAAATTAGATTAAGTTTGGGATTATGGTT
GTTGCAGGTGAAGTTGCGTGGGACCAACTTTCCTCCAGACGGTCCGGTTATGCAAAAGAAAACGATGG
GATGGGAAGCCAGCACCGAAAGACTTTATCCAGAGGATGTGGTGTTGAAGGGTGACATTAAGATGGCC
CTTAGGCTCAAAGATGGTGGAGATACTTGGCCGACTTCAAGACCACTTACCGTGCTAAAAAGCCAGT
GCAGATGCCAGGCGCATTCAATATCGACCGTAAGCTGGACATCACCTCGCATAATGAGGACTACACAG
TCGTGGAACAATACGAGCGTTCTGTCGCTCGTCATTCTACCGGTGGATCTGGTGGTTCTATGGACTCT
ACAGAAGCGGTTATCAAAGAATTCATGCGATTCAAGGTATTGAATCGTTTATTGCTTCTGTGATGTTG
TTGTTGCATTTGTGGCCTTCGTTGAGTTTTATGATCTGATTTCGATTGATACAGTTGTTTTATGATTC
TGTGTTGTTGTTGATGTTTCAATTGTTGCGTAATTGATTCGTTTAGGAACTGAACTTTTACTGGTGAT
GATTTAGGTCCACATGGAAGGATCGATGAATGGCCACGAGTTCGAAATAGAAGGTGAAGGCGAAGGTC
GTCCATACGAGGGTACACAAACCGCAAAGCTTCGTGTGACCAAAGGTGGTCCATTGCCGTTTAGCTGG
GACATCTTGTCTCCACAGTTTATGTATGGGTCCCGTGCCTTCACAAAGCACCCGGCCGATATTCCAGA
TTATTGGAAGCAGTCATTCCCTGAAGGCTTCAAATGGGAACGTGTTATGAACTTCGAGGACGGCGGTG
CCGTGTCAGTTGCACAAGATACAAGTTTGGAGGACGGCACATTGATCTACAAAGTCAAGCTGAGAGGG
ACGAACTTCCCACCGGATGGGCCAGTCATGCAAAAGAAAACTATGGGCTGGGAAGCGTCTACCGAAAG
GTAACATTCCTTAGTTACCTTTCTTTTCTTTTTCCATCATAAGTTTATAGATTGTACATGCTTTGAGA
TTTTTCTTTGCAAACAATCTCAGGTTGTACCCTGAAGATGTGGTCCTCAAAGGTGACATCAAAATGGC
GCTCCGTCTGAAAGACGGCGGAAGATATTTGGCGGATTTTAAGACGACCTATCGAGCCAAAAAGCCGG
TCCAAATGCCTGGTGCCTTTAATATCGACAGAAAACTCGACATTACCTCTCATAACGAAGATTATACC
GTCGTCGAGCAGTATGAGCGTTCAGTTGCTAGGCATAGCACTGGTGGTAGTGGTGGATCTTGA

edCitrineT9 (exon, intron)

GTGAGCAAGGGCGAGGAGCTGTTCACCGGGGTGGTGCCCATCCTGGTCGAGCTGGACGGCGACGTAAA
CGGCCACAAGTTCAGCGTGTCCGGCGAGGGCGAGGGCGATGCCACCTACGGCAAGCTGACCCTGAAGT
TCATCTGCACCACCGGCAAGCTGCCCGTGCCCTGGCCCACCCTCGTGACCACCTTCGGCTACGGCCTG
ATGTGCTTCGCCCGCTACCCCGACCACATGAAGCAGCACGACTTCTTCAAGTCCGCCATGCCCGAAGG
CTACGTCCAGGAGCGCACCATCTTCTTCAAGGACGACGGCAACTACAAGACCCGCGCCGAGGTAACAT
TCCTTAGTTACCTTTCTTTTCTTTTTCCATCATCTATCAATTTCTTTGCGGAAATTTATTTGAAGCTG
TAGAGTTAAAATTGAGTCTTTTAAACTTTTGTAGGTGAAGTTCGAGGGCGACACCCTGGTGAACCGCA
TCGAGCTGAAGGGCATCGACTTCAAGGAGGACGGCAACATCCTGGGGCACAAGCTTGAGTACAACTAC
AACAGCCACAACGTCTATATCATGGCCGACAAGCAGAAGAACGGCATCAAGGTAAGTTGTTACTTATG
ATTGTTTTCCTCTCTGCTACATGTATTTTGTTGTTCATTTCTGTAAGATATAAGAATTGAGTTTTCCT
CTGATGATATTATTAGGTGAACTTCAAGATCCGCCACAACATCGAGGACGGCAGCGTGCAGCTCGCCG
ACCACTACCAGCAGAACACCCCCATCGGCGACGGCCCCGTGCTGCTGCCCGACAACCACTACCTGAGC
TACCAGTCCGCCCTGTTCAAAGACCCCAACGAGAAGCGCGATCACATGGTCCTGCTGGAGTTCCTGAC
CGCCGCCGGGATC

T7edCerulean (exon, intron)

CCTCTGGTGTCAAAAGGCGAGGAGCTCTTCACGGGCGTCGTACCCATACTCGTTGAGCTTGATGGAGA
TGTGAATGGACATAAGTTTAGTGTTAGCGGAGAGGGAGAAGGTAAATCCTGGTCCACACTTTTACGAT
AAAAACACAAGATTTTAAACTATGAACTGATCAATAATCATTCCTAAAAGACCACACTTTTGTTTTGT
TTCTAAAGTAATTTTTACTGTTATAGCAGGAGATGCAACCTATGGAAAATTGACGTTGAAATTTATAT
GTACTACTGGTAAGCTGCCAGTTCCCTGGCCGACCCTCGTCACTACACTGTCCTGGGGGGTCCAATGC
TTCGCTAGATATCCTGATCATATGAAGCAACACGACTTTTTCAAAAGCGCCATGCCCGAAGGGTACGT
TCAGGAACGAACGATTTTCTTCAAAGATGACGGAAACTATAAGACGAGGGCGGAAGTAAAGTTTGAGG
GAGACACCTTGGTAAATCGAATAGAATTGAAGGGAATCGACTTTAAGGAAGATGGGAATATACTAGGC
CATAAGCTAGAGTATAATGCGATCCACGGGAACGTGTATATTACCGCGGACAAACAAAAAAATGGTAT
TAAAGCTAATTTTGGCCTCAACTGTAATATCGAGGATGGATCTGTTCAGTTAGCTGACCACTATCAAC
AAAATACACCGATAGGTAAAGCAACTGTGTTTTAATCAATTTCTTGTCAGGATATATGGATTATAACT
TAATTTTTGAGAAATCTGTAGTATTTGGCGTGAAATGAGTTTGCTTTTTGGTTTCTCCCGTGTTATAG
GTGACGGTCCCGTACTACTGCCCGACAATCACTACCTTAGCACCCAGTCTGCGCTATTCAAAGATCCT
AACGAGAAGCGGGACCATATGGTTCTTTTGGAGTTCTTGACCGCCGCGGGGATTACTTTAGGTATGGA
TGAGTTATACAAG

**Supplementary Table 1: Expression constructs**

| Name | Components | Assembly enzyme | Level | Description |
|---|---|---|---|---|
| pTGA2278 | pICSL4723, pTGA2207, pTGA2208, pTGA2277, pICH41766 | BpiI | 2 | RedSeed pAtAct2-SV40-3×mCitrine_intronised-6×HA pAtUBQ10-SV40-tagBFP-3×FLAG |
| pTGA2279 | pICSL4723, pTGA2207, pTGA2209, pTGA2277, pICH41766 | BpiI | 2 | RedSeed pAtAct2-SV40-3×mCitrine-6×HA pAtUBQ10-SV40-tagBFP-3×FLAG |
| pTGA2280 | pICSL4723, pTGA2207, pTGA2210, pTGA2277, pICH41766 | BpiI | 2 | RedSeed pAtAct2-SV40-3×mScarlet3_intronised-6×HA pAtUBQ10-SV40-tagBFP-3×FLAG |
| pTGA2281 | pICSL4723, pTGA2207, pTGA2211, pTGA2277, pICH41766 | BpiI | 2 | RedSeed pAtAct2-SV40-3×mScarlet3-6×HA pAtUBQ10-SV40-tagBFP-3×FLAG |
| pTGA2552 | pICSL4723, pTGA2207, pTGA2465, pTGA2517, pICH41766 | BpiI | 2 | RedSeed pSCR-SV40-3×mCitrine_intronised-6×HA-tSCR pAtUBQ10-tagBFP-3×FLAG |
| pTGA2553 | pICSL4723, pTGA2207, pTGA2466, pTGA2517, pICH41766 | BpiI | 2 | RedSeed pSCR-SV40-3×mCitrine-6×HA-tSCR pAtUBQ10-tagBFP-3×FLAG |
| pTGA2554 | pICSL4723, pTGA2207, pTGA2467, pTGA2517, pICH41766 | BpiI | 2 | RedSeed-pSCR-SV40-3×mScarlet3_intronised-6×HA-tSCR pAtUBQ10-tagBFP-3×FLAG |
| pTGA2555 | pICSL4723, pTGA2207, pTGA2468, pTGA2517, pICH41766 | BpiI | 2 | RedSeed pSCR-SV40-3×mScarlet3-6×HA-tSCR pAtUBQ10-tagBFP-3×FLAG |
| pTGA2716 | pICSL4723, pTGA2207, pTGA2706, pICH41744 | BpiI | 2 | FastRed_pUBQ10-SV40-6×HA-ABACUS2 E141D_100n no introns-tUBQ10 |
| pTGA2717 | pICSL4723, pTGA2207, pTGA2707, pICH41744 | BpiI | 2 | FastRed_pUBQ10-SV40-6×HA-ABACUS2 R143S_400n no introns-tUBQ10 |
| pTGA2718 | pICSL4723, pTGA2207, pTGA2708, pICH41744 | BpiI | 2 | FastRed_pUBQ10-SV40-6×HA-ABACUS2 E141D_100n intronised-tUBQ10 |
| pTGA2719 | pICSL4723, pTGA2207, pTGA2709, pICH41744 | BpiI | 2 | FastRed_pUBQ10-SV40-6×HA-ABACUS2 R143S_400n intronised-tUBQ10 |

**Supplementary Table 2: Acceptor Plasmids**

| Name | Origin | Level | Description |
|---|---|---|---|
| pICSL4723 | Castel B., Tomlinson L., Locci F., Yang Y., Jones J.D.G. (2019) PLoS ONE https://doi.org/10.1371/journal.pone.0204778 | 2 | Level2 acceptor |
| pICH47802 | pICH47802 was a gift from Sylvestre Marillonnet (Addgene plasmid # 48007 ; http://n2t.net/addgene:48007 ; RRID:Addgene_48007) | 1 | level1R acceptor |
| pICH47742 | pICH47742 was a gift from Sylvestre Marillonnet (Addgene plasmid # 48001 ; http://n2t.net/addgene:48001 ; RRID:Addgene_48001) | 1 | Level1 acceptor. Position2. Forward orientation |
| pICH47822 | pICH47822 was a gift from Sylvestre Marillonnet (Addgene plasmid # 48009 ; http://n2t.net/addgene:48009 ; RRID:Addgene_48009) | 1 | Level1 acceptor. Position3. Reverse orientation |
| pICH47751 | pICH47751 was a gift from Sylvestre Marillonnet (Addgene plasmid # 48002 ; http://n2t.net/addgene:48002 ; RRID:Addgene_48002) | 1 | Level1 acceptor. Position3. Forward orientation |
| pICH41264 | pICH41264 was a gift from Sylvestre Marillonnet (Addgene plasmid # 47993 ; http://n2t.net/addgene:47993 ; RRID:Addgene_47993) | 0 | CDS2 acceptor |
| pICH41258 | pICH41258 was a gift from Sylvestre Marillonnet (Addgene plasmid # 47987 ; http://n2t.net/addgene:47987 ; RRID:Addgene_47987) | 0 | NT2 acceptor |
| pICH41308 | pICH41308 was a gift from Sylvestre Marillonnet (Addgene plasmid # 47998 ; http://n2t.net/addgene:47998 ; RRID:Addgene_47998) | 0 | CDS1 acceptor |
| pAGM1301 | pAGM1301 was a gift from Sylvestre Marillonnet (Addgene plasmid # 47989 ; http://n2t.net/addgene:47989 ; RRID:Addgene_47989) | 0 | CT acceptor |
| pAGM1299 | pAGM1299 was a gift from Sylvestre Marillonnet (Addgene plasmid # 47988 ; http://n2t.net/addgene:47988 ; RRID:Addgene_47988) | 0 | CDS2ns acceptor |
| pAGM1311 | pAGM1311 was a gift from Sylvestre Marillonnet (Addgene plasmid # 47983 ; http://n2t.net/addgene:47983 ; RRID:Addgene_47983) | -1 | Level -1 acceptor |
| p4L1r pDONR | Andersen T.G., Molina D., Kilian J., Franke R.B., Ragni L., Geldner N. (2021) Current Biology; doi.org/10.1016/j.cub.2020.11.070 | 0 | modified acceptor for gateway and golden gate |
| pR2-L3 | Invitrogen | GW | modified acceptor for gateway and golden gate |

**Supplementary Table 3: Level1 Plasmids**

| Name | Components / Origin | enzyme | Module | Description |
|---|---|---|---|---|
| pTGA2207 | pICH47802, pICSL70008 | BsaI | 1R | FastRed |
| pTGA2208 | pICH47742, pICH87644, pTGA2201, pTGA2206, pICSL50009, pICH44300 | BsaI | 2F | pAtAct2-SV40-3×mCitrine_intronised-linker-6×HA-tAtAct2 |
| pTGA2209 | pICH47742, pICH87644, pTGA2202, pTGA2206, pICSL50009, pICH44300 | BsaI | 2F | pAtAct2-SV40-3×mCitrine-linker-6×HA-tAtAct2 |
| pTGA2210 | pICH47742, pICH87644, pTGA2203, pTGA2206, pICSL50009, pICH44300 | BsaI | 2F | pAtAct2-SV40-3×mScarlet3_intronised-linker-6×HA-tAtAct2 |
| pTGA2211 | pICH47742, pICH87644, pTGA2203, pTGA2206, pICSL50009, pICH44300 | BsaI | 2F | pAtAct2-SV40-3×mScarlet-linker-6×HA-tAtAct2 |
| pTGA2277 | pICH47822, pTGA1892, pTGA2227, pTGA2228, pICSL50007, pTGA1894 | BsaI | 3R | pAtUBQ10-SV40-tagBFP-linker-3×FLAG-tAtUBQ10 |
| pTGA2465 | pICH47742, pTGA1039, pTGA2201, pTGA2206, pICSL50009, pTGA2460 | BsaI | 2F | pSCR-SV40-3×mCitrine_intronised-linker-6×HA-tSCR |
| pTGA2466 | pICH47742, pTGA1039, pTGA2202, pTGA2206, pICSL50009, pTGA2460 | BsaI | 2F | pSCR-SV40-3×mCitrine-linker-6×HA-tSCR |
| pTGA2467 | pICH47742, pTGA1039, pTGA2203, pTGA2206, pICSL50009, pTGA2460 | BsaI | 2F | pSCR-SV40-3×mScarlet3_intronised-linker-6×HA-tSCR |
| pTGA2468 | pICH47742, pTGA1039, pTGA2204, pTGA2206, pICSL50009, pTGA2460 | BsaI | 2F | pSCR-SV40-3×mScarlet-linker-6×HA-tSCR |
| pTGA2517 | pICH47751, pTGA1892, pTGA2227, pTGA2228, pICSL50007, pTGA1894 | BsaI | 3F | pAtUBQ10-SV40-tagBFP-linker-3×FLAG-tAtUBQ10 |
| pTGA2706 | pICH47772, pTGA1892, pTGA2651, pTGA2659, pTGA1894 | BsaI | 2F | pUBQ10-SV40-6×HA-ABACUS2 E141D_100n no introns-tUBQ10 |
| pTGA2707 | pICH47772, pTGA1892, pTGA2651, pTGA2660, pTGA1894 | BsaI | 2F | pUBQ10-SV40-6×HA-ABACUS2 R143S_400n no introns-tUBQ10 |
| pTGA2708 | pICH47772, pTGA1892, pTGA2651, pTGA2661, pTGA1894 | BsaI | 2F | pUBQ10-SV40-6×HA-ABACUS2 E141D_100n intronised-tUBQ10 |
| pTGA2709 | pICH47772, pTGA1892, pTGA2651, pTGA2662, pTGA1894 | BsaI | 2F | pUBQ10-SV40-6×HA-ABACUS2 R143S_400n intronised-tUBQ10 |
| pICH41744 | pICH41744 was a gift from Sylvestre Marillonnet (Addgene plasmid # 48017 ; | BsaI | ELE2 | End-link 2 for assembling 2 level one part into a level 2 acceptor |

| | | | | |
|---|---|---|---|---|
| | http://n2t.net/addgene:48017 ; RRID:Addgene_48017) | | | |
| pICH41766 | pICH41766 was a gift from Sylvestre Marillonnet (Addgene plasmid # 48018 ; http://n2t.net/addgene:48018 ; RRID:Addgene_48018) | BsaI | ELE3 | End-link 3 for assembling 3 level one part into a level 2 acceptor |

**Supplementary Table 4: Level0 Plasmids**

| Name | Components / Origin | Enzym e | Module | Description |
|---|---|---|---|---|
| pICSL7000 8 | pICSL70008 was a gift from Nicola Patron (Addgene plasmid # 50336 ; http://n2t.net/addgene:50336 ; RRID:Addgene_50336) | | | FastRed |
| pTGA2201 | pICH41258, gSAS216AN, gSAS216B,gSAS 216C,gSAS 216D, gSAS217Enone | BpiI | NT2 | SV40-linker-3×mCitrine_intronised |
| pTGA2202 | pICH41258, gSAS216AN, gSAS218Ba, gSAS218Bb, gSAS217Enone | BpiI | NT2 | SV40-linker-3×mCitrine |
| pTGA2203 | pICH41258, gSAS222AN, gSAS222B, gSAS222C, gSAS223Dnone | BpiI | NT2 | SV40-linker-3×mScarlet3_intronised |
| pTGA2204 | pICH41258, gSAS222AN, gSAS224Ba, gSAS224Bb, gSAS223Dnone | BpiI | NT2 | SV40-linker-3×Scarlet3 |
| pTGA2190 | pICH41258, gSAS217A, gSAS216B, gSAS216C, gSAS216D, gSAS217Enone | BpiI | NT2 | 3×mCitrine_intronised |
| pTGA2192 | pICH41258, gSAS217A, gSAS218Ba, gSAS218Bb, gSAS217Enone | BpiI | NT2 | 3×mCitrine |
| pTGA2196 | pICH41258, gSAS223A, gSAS222B, gSAS222C, gSAS223Dnone | BpiI | NT2 | 3×mScarlet3_intronised |
| pTGA2198 | pICH41258, gSAS223A, gSAS224Ba, gSAS224Bb, gSAS223Dnone | BpiI | NT2 | 3×mScarlet3 |
| pTGA2189 | pICH41308, gSAS216AN, gSAS216B, gSAS216C, gSAS216D, gSAS216Estop | BpiI | CDS1 | SV40-linker-3×mCitrine_intronised |

| | | | | |
|---|---|---|---|---|
| pTGA2191 | pICH41308, gSAS216AN, gSAS218Ba, gSAS218Bb, gSAS216Estop | BpiI | CDS1 | SV40-3×mCitrine-STOP |
| pTGA2195 | pICH41308, gSAS222AN, gSAS222B, gSAS222C, gSAS222Dstop | BpiI | CDS1 | SV40-linker-3×mScarlet3_intronised |
| pTGA2197 | pICH41308, gSAS222AN, gSAS224Ba, gSAS224Bb, gSAS222Dstop | BpiI | CDS1 | SV40-linker-3×mScarlet3 |
| pTGA2193 | pAGM1301, gSAS220A, gSAS216B, gSAS216C, gSAS216D, gSAS216Estop | BpiI | CT | 3×mCitrine_intronised |
| pTGA2194 | pAGM1301, gSAS220A, gSAS218Ba, gSAS218Bb, gSAS216Estop | BpiI | CT | 3×mCitrine-STOP |
| pTGA2199 | pAGM1301, gSAS226A, gSAS226B, gSAS226C, gSAS222Dstop | BpiI | CT | 3×mScarlet3_intronised |
| pTGA2200 | pAGM1301,gSAS226A, gSAS224Ba, gSAS224Bb, gSAS223Dnone | BpiI | CT | 3×mScarlet3 |
| pTGA2205 | pICH41258, gSAS232A | BpiI | NT2 | tagBFP |
| pTGA2206 | pAGM1299, gSAS223A | BpiI | CDS2ns | flexible linker |
| pICSL50009 | pICSL50009 was a gift from Nicola Patron (Addgene plasmid # 50309 ; http://n2t.net/addgene:50309 ; RRID:Addgene_50309) | | CT | 6×HA tag (6x Human influenza hemagglutinin) |
| pICH87644 | pICH87644 was a gift from Sylvestre Marillonnet & Nicola Patron (Addgene plasmid # 50274 ; http://n2t.net/addgene:50274 ; RRID:Addgene_50274) | | P | pAtAct2 and 5'UTR, omega (Tobacco Mosaic Virus) |
| pICH44300 | pICH44300 was a gift from Sylvestre Marillonnet & Nicola Patron (Addgene plasmid # 50340 ; http://n2t.net/addgene:50340 ; RRID:Addgene_50340) | | 3UT | tAtAct2 |
| pTGA1892 | p4L1r pDONR, part1892 | infusion | P | pUBQ10 |
| pTGA2227 | pICH41258, SAS257 | BpiI | NT2 | SV40-linker |
| pTGA2228 | pAGM1299, SAS258 | BpiI | CDS2ns | tagBFP-linker |
| pICSL50007 | pICSL50007 was a gift from Nicola Patron (Addgene plasmid # 50308 ; http://n2t.net/addgene:50308 ; RRID:Addgene_50308) | | CT | 3×FLAG tag (3x FLAG octapeptide) |
| pTGA1894 | pR2-L3; part1894 | infusion | 3UT | UBQ10 3'UTR |
| pTGA1039 | p4L1r pDONR, part1039A, part1039B, part1039C, part1039D Andersen et al., (2018) Nature 10.1038/nature25976 | infusion | P | pSCR |

| | | | | |
|---|---|---|---|---|
| pTGA2460 | pICH41276, SAS320 | BpiI | 3UT | 3'UTR tSCR |
| pTGA2651 | part457A, part457B, part457C, part457D | BpiI | NT2 | SV40-linker-6×HA-linker |
| pTGA2659 | pICH41264, pTGA2648, pTGA2649, pTGA2650, Jim09, Jim11 | BpiI | CDS2 | ABACUS2 E141D_100n no introns |
| pTGA2660 | pICH41264, pTGA2648, pTGA2649, pTGA2650, Jim10, Jim11 | BpiI | CDS2 | ABACUS2 R143S_400n no introns |
| pTGA2661 | pICH41264, pTGA2646, pTGA2649, pTGA2649, Jim09, pTGA2647 | BpiI | CDS2 | ABACUS2 E141D_100n intronised |
| pTGA2662 | pICH41264, pTGA2646, pTGA2649, pTGA2649, Jim10, pTGA2647 | BpiI | CDS2 | ABACUS2 R143S_400n intronised |

**Supplementary Table 5: Level -1 modules**

| Name | Components / Origin | Enzyme | Module | Description |
|---|---|---|---|---|
| pTGA2646 | pAGM1311, gSAS452 | BsaI | -1 | AGGT-TCCG-edCIT t9 intronised |
| pTGA2647 | pAGM1311, gSAS453 | BsaI | -1 | CCTC-GCTT-PL-edCer3 intronised |
| pTGA2648 | pAGM1311, gSAS454 | BsaI | -1 | AGGT-TCCG-edCIT t9 |
| pTGA2649 | pAGM1311, SAS455A, SAS455B, SAS455C | BsaI | -1 | TCCG-TGGA-ABI1ait |
| pTGA2650 | pAGM1311, SAS456A, SAS456B, SAS456C | BsaI | -1 | TGGA-GGAG-L52 |
| PYL1 (100n) | was a gift from James Rowe (Rowe et al., 2023) | | -1 | GGAG-CCTC-PYL1 H87P S112A E141D A140V (100n) |
| PYL1 (400n) | was a gift from James Rowe (Rowe et al., 2023) | | -1 | GGAG-CCTC-PYL1 H87P S112A R143S A140V (400n) |
| edCer3 | was a gift from James Rowe (Rowe et al., 2023) | | -1 | CCTC-GCTT-edCer3 |

**Supplementary Table 6: PCR products, templates, and primers**

| Name | PCR configuration |
|---|---|
| SAS257 | primer template: b369; primer: b365 + b366 = 85 bp |
| SAS258 | template: pTGA2205; primer: b370 + b371 = 743 bp |
| SAS320 | template: Col-0 gDNA; primer: b421 + b422 =283 bp |
| SAS454A | primer template: b474; primer: b472 + b473 = 98 bp |
| SAS454B | primer template: b475; primer: b472 + b473 = 87 bp |
| SAS454C | primer template: b476; primer: b472 + b473 = 92 bp |
| SAS456A | primer template: b477; primer: b473 = 100 bp |
| SAS456B | primer template: b478; primer: b473 = 95 bp |
| SAS456C | primer template: b479; primer: b473 = 99 bp |
| part457A | primer template: b468; primer: b365 + b366 = 101 bp |
| part457B | primer template: b469; primer: b365 + b366 = 106 bp |
| part457C | primer template: b470; primer: b365 + b366 = 97 bp |
| part457D | primer template: 471; primter: b365 + b366 = 114 bp |
| part1892 | template: Col-0 gDNA; primer: t001 + t002 = 2038 bp |
| part1894 | template: Col-0 gDNA; primer: t003 + t004 = 435 bp |
| part1039A | template: Col-0 gDNA; primer: t005 + t006 = 1782 bp |
| part1039B | template: Col-0 gDNA; primer: t007 + t008 = 219 bp |
| part1039C | template: Col-0 gDNA; primer: t009 + t010 = 182 bp |
| part1039D | template: Col-0 gDNA; primer: t011 + t012 = 206 bp |

**Supplementary Table 7: Primers**

| Name | Sequence |
|---|---|
| b365 | GATCGGTTGTGAAGACTT |
| b366 | TCGTACGCTCGAAGACTT |
| b369 | GATCGGTTGTGAAGACTTAATGATGCCAAAGAAGAAGAGGAAGGTTGGCGGCGGAGG TAGCGGAGGTAAGTCTTCGAGCGTACGA |

| | |
|---|---|
| b370 | CGTGAAGACTAAGGTTCTGAACTTATTAAGGAGAA |
| b371 | AGCGAAGACATCGAACCGCTACCTCCGCCGCCATTGAGCTTGTGTCCAAG |
| b421 | TGAGAAGACTAGCTTTTTTCTTCTCCTTTTTCACAAAC |
| b422 | AGAGAAGACTTAGCGTATTTGTAAATGATTTTTCAG |
| b468 | GATCGGTTGTGAAGACTTAATGCCAAAGAAGAAGAGGAAGGTTGGCGGCGGAGGTAG CGGTGGATATCCTTATGATGTTCCTAAGTCTTCGAGCGTACGA |
| b469 | GATCGGTTGTGAAGACTTTCCTGATTATGCTGGATATCCATATGATGTTCCAGATTATGCT GGTCTTTACCCTTACGATGTTCCTGAAAGTCTTCGAGCGTACGA |
| b470 | GATCGGTTGTGAAGACTTCTGATTATGCTACTAGAGCTGCTTACCCTTACGATGTTCCTG ATTATGCTGGTTACCCATAAGTCTTCGAGCGTACGA |
| b471 | GATCGGTTGTGAAGACTTCCATACGATGTTCCTGATTACGCTGGACTCTACCCTTACGA TGTTCCTGATTATGCTGGAGGAGGAGGATCTGGAGGTAAGTCTTCGAGCGTACGA |
| b472 | GTATAGTGCATTGGTCTCA |
| b473 | GATTAGTGCATTGGTCTCA |
| b474 | GTATAGTGCATTGGTCTCAACATTCCGCATAAACCGGATAGAGAAGATGAAGCTGCGAG GATTGAAGCCGCAGGAGGGAAAGTGTGAGACCAATGCACTAATC |
| b475 | GTATAGTGCATTGGTCTCAAGTGATTCAGTGGAATGGAGCTCGTGTTTTCGGTGTTCTCG CCATGTCGTGAGACCAATGCACTAATC |
| b476 | GTATAGTGCATTGGTCTCAGTCGAGATCCATTGGCGATAGATACTTGAAACCATCCATCAT TCCTGATGGTGGATTGTTGAGACCAATGCACTAATC |
| b477 | GTATAGTGCATTGGTCTCAACATTGGAGCAAGTGGCGCGCCTGGTCCTGGAGGTGCAGG TCCAGGAGGTGCAGGACCAGGTTGAGACCAATGCACTAATC |
| b478 | GTATAGTGCATTGGTCTCAAGGTGGAGCTGGTCCTGGTGGAGCTGGACCTGGAGGTGCA GGACCTGGAGGAGCTGGTGAGACCAATGCACTAATC |
| b479 | GTATAGTGCATTGGTCTCACTGGTCCTGGTGGTGCTGGACCTGGTGGAGCTGGGGCCGG CCGAAGTTCTGGAGGAGTTGTTGAGACCAATGCACTAATC |
| t001 | ATAGAAAAGTTGGATGGTCTCAGGAGAGTCTAGCTCAACAGAGCTTTTAACC |
| t002 | ACAAACTTGCGGATCGGTCTCACATTCTGTTAATCAGAAAAACTCAGATTAATCGAC |
| t003 | CAAAGTGGTAGGTCTCAGCTTATCTCGTCTCTGTTATGCTTAAGAAG |
| t004 | AAAGTTGTGGTCTCAAGCGGCAACAACGAATGTTCATCATACTCA |
| t005 | TGTACAAGAAAGCTGGCTTATCTGTGGTCTCAGGAGAGACATACTCCAGACTCTGC |
| t006 | GTGTGTTTGACGTGTTCCGTTTGCC |

| | |
|---|---|
| t007 | GGCAAACGGAACACGTCAAACACAC |
| t008 | GCCGTATCTAAGTCGTGTTCCACC |
| t009 | GGTGGAACACGACTTAGATACGGC |
| t010 | GATTCTTAAATTACTTGATCGTGTTCACGATCTCC |
| t011 | GGAGATCGTGAACACGATCAAGTAATTTAAGAATC |
| t012 | TTGTACAAAAAAGCTGGGTTTCGTGGTCTCTCATTGGGAGATTGAAGGGTTGTTG |

**Supplementary Table 8: Synthesised gBlocks**

| Name | Sequence |
|---|---|
| gSAS 216AN | TCTGCTATTAAGCCTGATATGAAGATTAAATTTGAAGACTTAATGCCAAAGAAGAAGAGGAAGGTTGGCGGCGGAGGTAGCATGGTGA GCAAGGGCGAAGAGTTGTTCACTGGTGTTGTTCCTATCCTCGTTGAGCTTGACGGTGATGTGAACGGGCATAAGTTCAAGTCTTCAAAC AGAATGGAGGGAAACGTTAACGG |
| gSAS 216B | GAAGACTTGTTCTCCGTTTCTGGTGAAGGTAACATTCCTTAGTTACCTTTCTTTTCTTTTTCCATCATCTATCAATTTCTTTGCGGAAATTT ATTTGAAGCTGTAGAGTTAAAATTGAGTCTTTTAAACTTTTGTAGGTGAGGGAGATGCTACTTACGGAAAGCTCACCCTCAAGTTCATCT GTACCACTGGAAAGCTCCCTGTGCCTTGGCCTACTCTCGTTACTACTTTCGGATACGGGCTCATGTGCTTCGCTAGATACCCTGATCAT ATGAAGCAGCACGACTTCTTCAAGAGCGCTATGCCTGAGGGATACGTGCAAGAGAGAACCATTTTCTTCAAGGACGACGGGAACTACA AGACCAGAGCTGAGGTTAAGTTCGAAGGTGACACCCTCGTGAACAGGATCGAGCTTAAGGGCATCGACTTCAAAGAGGACGGAAACAT CCTCGGACACAAGCTCGAGTACAACTACAACAGCCACAACGTGTACATCATGGCCGACAAGCAGAAGAACGGCATCAAGGTAAGTTGT TACTTATGATTGTTTTCCTCTCTGCTACATGTATTTTGTTGTTCATTTCTGTAAGATATAAGAATTGAGTTTTCCTCTGATGATATTATTAG GTCAACTTCAAGATCAGGCACAACATCGAGGACGGATCTGTTCAGCTCGCTGATCATTACCAGCAGAACACCCCTATTGGAGATGGAC CTGTTCTTCTCCCTGACAACCACTACCTCAGCTACCAGTCTAAGCTCAGCAAGGACCCTAACGAGAAGAGGGATCATATGGTGCTCCTC GAGTTCGTTACTGCTGCTGGAATCACTCTCGGAATGGACGAGCTTTACAAAGGCGGTGGTGGATCCATGGTGTCCAAGGGTGAAGAAC TTTTCACAGGTGTGGTGCCGATCCTTGTGGAACTCGATGGTGATGTCAATGGCCACAAGTTCAGCGTGAGCGGAGAAGGCGAAGGTAA ATCCTAAGTCTTC |

| | |
|---|---|
| gSAS 216C | GAAGACTTTCCTGGTCCACACTTTTACGATAAAAACACAAGATTTTAAACTATGAACTGATCAATAATCATTCCTAAAAGACCACACTTTT<br>GTTTTGTTTCTAAAGTAATTTTTACTGTTATAACAGGTGATGCAACATATGGTAAGTTGACGCTGAAGTTTATCTGTACGACCGGGAAGTT<br>GCCTGTTCCATGGCCAACACTTGTGACCACATTCGGCTACGGATTGATGTGTTTCGCTCGTTACCCAGACCACATGAAGCAACATGATT<br>TCTTTAAGTCCGCCATGCCAGAGGGCTATGTCCAAGAAAGGACGATCTTTTTCAAGGATGATGGCAATTATAAGACCCGTGCCGAAGTG<br>AAATTCGAGGGCGATACTCTTGTGAACCGTATCGAGTTGAAAGGTAAGTTCTGCATTTGGTTATGCTCCTTGCATTTTAGGTGTTCGTCG<br>CACTTCCATTTCCATGAATAGCTAAGATTTTTTTTCTCTGCATTCATTCTTCTTGCCTCAGTTCTAACTGTTTGTGGTATTTTTGTTTTAATT<br>ATTGCTACAGGTATTGATTTTAAAGAGGATGGGAACATTTTGGGGCACAAGTTGGAGTATAATTACAACTCGCATAACGTCTACATTATG<br>GCGGATAAGCAAAAGAATGGTATCAAAGTGAATTTTAAGATCCGTCATAACATCGAAGATGGCAGCGTGCAGTTGGCAGATCACTACCA<br>ACAAAACACGCCGATCGGTGATGGTCCTGTGCTTCTTCCAGATAACCATTACCTGTCCTACCAGAGCAAGCTCTCTAAGGATCCGAATG<br>AGAAGCGTGATCACATGGTCCTTTTGGAGTTTGTGACCGCTGCTGGTATTACCCTTGGCATGGATGAACTCTACAAAGGCGGCGGAGG<br>TAGCATGGTTTCTAAAGGTAAAGCAACTGTGTTTTAATCAATTTCTTGTCAGGATATATGGATTATAACTTAATTTTTGAGAAAAAGTCTTC |
| gSAS 216D | GAAGACTTGAAATCTGTAGTATTTGGCGTGAAATGAGTTTGCTTTTTGGTTTCTCCCGTGTTATAGGTGAGGAACTCTTTACCGGGGTCG<br>TGCCTATTCTTGTCGAGCTTGATGGGGACGTTAACGGCCACAAATTCTCGGTGTCAGGTGAAGGTGAAGGCGACGCTACCTATGGGAA<br>GCTCACTTTGAAATTCATATGTACTACGGGCAAGCTGCCGGTTCCTTGGCCGACTTTGGTTACAACATTTGGCTATGGCCTGATGTGCT<br>TTGCCAGGTATCCGGACCATATGAAGCAGCATGATTTCTTCAAAAGTGCCATGCCGGAAGGTTATGTGCAAGAGCGTACGATATTCTTT<br>AAAGATGACGGTAATTACAAGACGAGGGCCGAAGTCAAGTTTGAAGGGGATACACTGGTCAACCGAATCGAACTCAAGGGAATAGATT<br>TCAAAGAAGATGGTAATATCCTGGGGCATAAGCTAGAGTATAACTATAATTCCCACAATGTCTATATCATGGCTGACAAACAAAAGAACG<br>GTATTAAGGTAAAGTTTCCAACTTTCCTTTACCATATCAAACTAAAGTTCGAAACTTTTTATTTGATCAACTTCAAGGCCACCCGATCTTTC<br>TATTCCTGATTAATTTGTGATGAATCCATATTGACTTTTGATGGTTACGCAGGTGAACTTCAAAATCCGACATAATATTGAGGACGGCAG<br>CGTCCAGCTTGCTGACCACTATCAACAGAATACCCCAATCGGTGACGGACCAGTTCTCTTGCCTGATAATCACTACTTGTCGTACCAAA<br>GTCTTC |
| gSAS 216Esto | CAGAATGGAGGGAAACGTTAACGGTTTGAAGACTTACCAGTCCAAGTTGTCGAAAGATCCTAACGAAAAGCGTGACCACATGGTGTTG<br>CTCGAGTTTGTCACAGCAGCAGGTATCACGCTCGGTATGGATGAGTTGTATAAGTGAGCTTAAGTCTTCAAATCATTTCGTTATCGATG<br>GAGAGGGTACAGGTAAACCTTTC |
| gSAS 217A | TCATTTCGTTATCGATGGAGAGGGTACAGGTAAACCTTTCTTTGAAGACTTAATGGTGAGCAAGGGCGAAGAGTTGTTCACTGGTGTTG<br>TTCCTATCCTCGTTGAGCTTGACGGTGATGTGAACGGGCATAAGTTCAAGTCTTCAAACTTACAACTGCTTTCCATTATGGAAATAGAGT<br>TTTTGCTAAATACCCTGATAA |
| gSAS 217Enon | TGCTTTCCATTATGGAAATAGAGTTTTTGCTAAATACCCTGATAATTTGAAGACTTACCAGTCCAAGTTGTCGAAAGATCCTAACGAAAAG<br>CGTGACCACATGGTGTTGCTCGAGTTTGTCACAGCAGCAGGTATCACGCTCGGTATGGATGAGTTGTATAAGGGAGGTAAGTCTTCAA<br>ATACCCTGATAACATCCAAGA |

| | |
|---|---|
| gSAS 218Ba | GAAGACTTGTTCTCCGTTTCTGGTGAAGGTGAGGGAGATGCTACTTACGGAAAGCTCACCCTCAAGTTCATCTGTACCACTGGAAAGCT<br>CCCTGTGCCTTGGCCTACTCTCGTTACTACTTTCGGATACGGGCTCATGTGCTTCGCTAGATACCCTGATCATATGAAGCAGCACGACT<br>TCTTCAAGAGCGCTATGCCTGAGGGATACGTGCAAGAGAGAACCATTTTCTTCAAGGACGACGGGAACTACAAGACCAGAGCTGAGGT<br>TAAGTTCGAAGGTGACACCCTCGTGAACAGGATCGAGCTTAAGGGCATCGACTTCAAAGAGGACGGAAACATCCTCGGACACAAGCTC<br>GAGTACAACTACAACAGCCACAACGTGTACATCATGGCCGACAAGCAGAAGAACGGCATCAAGGTCAACTTCAAGATCAGGCACAACA<br>TCGAGGACGGATCTGTTCAGCTCGCTGATCATTACCAGCAGAACACCCCTATTGGAGATGGACCTGTTCTTCTCCCTGACAACCACTAC<br>CTCAGCTACCAGTCTAAGCTCAGCAAGGACCCTAACGAGAAGAGGGATCATATGGTGCTCCTCGAGTTCGTTACTGCTGCTGGAATCA<br>CTCTCGGAATGGACGAGCTTTACAAAGGCGGTGGTGGATCCATGGTGTCCAAGGGTGAAGAACTTTTCACAGGTGTGGTGCCGATCCT<br>TGTGGAACTCGATGGTGATGTCAATGGCCACAAGTTCAGCGTGAGCGGAGAAGGCGAAGGTGATGCAACATATGGTAAGTTGACGCT<br>GAAGTTTATCTGTACGACCGGGAAGTTGCCTGTTCCATGGCCAACACTTGTGACCACATTCGGCTACGGATTGATGTGTTTCGCTCGTT<br>ACCCAGACCACATGAAGCAACATGATTTCTTTAAGTCCGCCATGCCAGAGGGCTATGTCCAAGAAAGGACGATCTTTTTCAAGGATGAT<br>GGCAATTATAAGTCTTC |
| gSAS 218Bb | GAAGACTTTTATAAGACCCGTGCCGAAGTGAAATTCGAGGGCGATACTCTTGTGAACCGTATCGAGTTGAAAGGTATTGATTTTAAAGA<br>GGATGGGAACATTTTGGGGCACAAGTTGGAGTATAATTACAACTCGCATAACGTCTACATTATGGCGGATAAGCAAAAGAATGGTATCA<br>AAGTGAATTTTAAGATCCGTCATAACATCGAAGATGGCAGCGTGCAGTTGGCAGATCACTACCAACAAAACACGCCGATCGGTGATGGT<br>CCTGTGCTTCTTCCAGATAACCATTACCTGTCCTACCAGAGCAAGCTCTCTAAGGATCCGAATGAGAAGCGTGATCACATGGTCCTTTT<br>GGAGTTTGTGACCGCTGCTGGTATTACCCTTGGCATGGATGAACTCTACAAAGGCGGCGGAGGTAGCATGGTTTCTAAAGGTGAGGAA<br>CTCTTTACCGGGGTCGTGCCTATTCTTGTCGAGCTTGATGGGGACGTTAACGGCCACAAATTCTCGGTGTCAGGTGAAGGTGAAGGCG<br>ACGCTACCTATGGGAAGCTCACTTTGAAATTCATATGTACTACGGGCAAGCTGCCGGTTCCTTGGCCGACTTTGGTTACAACATTTGGC<br>TATGGCCTGATGTGCTTTGCCAGGTATCCGGACCATATGAAGCAGCATGATTTCTTCAAAAGTGCCATGCCGGAAGGTTATGTGCAAGA<br>GCGTACGATATTCTTTAAAGATGACGGTAATTACAAGACGAGGGCCGAAGTCAAGTTTGAAGGGGATACACTGGTCAACCGAATCGAA<br>CTCAAGGGAATAGATTTCAAAGAAGATGGTAATATCCTGGGGCATAAGCTAGAGTATAACTATAATTCCCACAATGTCTATATCATGGCT<br>GACAAACAAAAGAACGGTATTAAGGTGAACTTCAAAATCCGACATAATATTGAGGACGGCAGCGTCCAGCTTGCTGACCACTATCAACA<br>GAATACCCCAATCGGTGACGGACCAGTTCTCTTGCCTGATAATCACTACTTGTCGTACCAAAGTCTTC |
| gSAS 220A | TACCCTGATAACATCCAAGATTTGAAGACTTTTCGATGGTGAGCAAGGGCGAAGAGTTGTTCACTGGTGTTGTTCCTATCCTCGTTGAG<br>CTTGACGGTGATGTGAACGGGCATAAGTTCAAGTCTTCAAATTACAACTGCTTTCCATTATGGAAATAGAGTTTTTGCTAAATACCCTGA<br>TAACATCCAAGATTATTTCAA |
| gSAS 222AN | CGGAACCAATTTTCCTGCTAATGGTCCAGTCATTTGAAGACTTAATGCCAAAGAAGAAGAGGAAGGTTGGCGGCGGAGGTAGCATGGA<br>TTCTACCGAGGCCGTGATCAAAGAATTCATGAGGTTCAAGTCTTCAAACTGCTTTCCATTATGGAAATAGAGTTTTTGCTAAATACCCTG<br>ATAACATCCAAGATTATTTCAA |

| | |
|---|---|
| gSAS 222B | GAAGACTTGTTCAAGGTGCACATGGAAGGCAGCATGAACGGACATGAGTTCGAGATCGAAGGTGAAGGTGAGGGTAGACCTTACGAG<br>GGAACTCAGACTGCTAAGCTCAGAGTGACAAAAGGTCTGTCTTTCCTATTTCATATGTTTAATCCTAGGAATTTGATCAATTGATTGTATG<br>TATGTCGATCCCAAGACTTTCTTGTTCACTTATATCTTAACTCTCTCTTTGCTGTTTCTTGCAGGTGGACCGCTTCCGTTCTCTTGGGATA<br>TCCTTTCACCGCAGTTCATGTACGGCTCTAGGGCTTTTACTAAGCACCCTGCTGATATCCCGGACTACTGGAAGCAATCTTTCCCAGAG<br>GGATTCAAGTGGGAGCGTGTGATGAATTTCGAGGATGGTGGTGCTGTTTCTGTGGCTCAGGATACTTCTCTTGAGGACGGAACCCTCA<br>TCTACAAGGTAATATTTTGATAGAGATTATTCTATTCAGCATTGAAGCATCGTTGCTACAAATGGTACCATGAACTTTTTGTTTACTTCTCT<br>CTACCTCTATAGGTTAAGCTCCGTGGAACTAACTTCCCTCCTGATGGACCTGTGATGCAGAAAAAGACCATGGGCTGGGAAGCTTCTAC<br>TGAGAGACTTTACCCTGAGGACGTTGTGCTCAAGGGCGATATCAAGATGGCTCTCAGACTTAAGGACGGTGGACGTTACCTCGCTGAT<br>TTCAAGACTACCTACAGGGCCAAGAAACCTGTGCAAATGCCTGGGGCTTTCAACATCGATAGGAAGCTCGATATCACCAGCCACAACG<br>AGGATTACACTGTGGTTGAGCAGTACGAGAGATCTGTGGCTAGACACTCTACTGGTGGAAGCGGAGGATCTATGGATAGCACTGAGGC<br>TGTCATCAAAGAATTCATGCGTTTCAAAGTTCATATGGAAGGTAAAATATTGGATGCCAGACGATATTCTTTCTTTTGATTTGTAACTTTTT<br>CCTGTCAAGGTCGATAAATTTTATTTTTTTTGGTAAAAGGTCGATAATTTTTTTTTGGAGCCATTATGTAATTTTCCTAATTAACTGAACCA<br>AAATTATACAAACCAGGTTCCATGAATGGGCAAAGTCTTC |
| gSAS 222C | GAAGACTTGGCACGAATTTGAAATTGAAGGCGAAGGCGAGGGACGTCCGTATGAAGGTACTCAAACAGCTAAGCTGAGGGTTACCAAA<br>GGCGGTCCTCTTCCTTTTTCCTGGGATATCTTGAGCCCTCAATTCATGTATGGCAGCAGGGCCTTCACCAAACATCCAGCAGATATCCC<br>TGATTATTGGAAACAAAGCTTCCCGGAAGGGTTTAAGTGGGAGAGAGTCATGAACTTCGAAGATGGCGGAGCAGTGAGTGTTGCTCAA<br>GACACTTCTTTGGAGGATGGGACTCTGATCTATAAGGTAAGAAAATGATTCCTCATAGAGTAAACAAATGGTAGATGAAAATTAGATTAA<br>GTTTGGGATTATGGTTGTTGCAGGTGAAGTTGCGTGGGACCAACTTTCCTCCAGACGGTCCGGTTATGCAAAAGAAAACGATGGGATG<br>GGAAGCCAGCACCGAAAGACTTTATCCAGAGGATGTGGTGTTGAAGGGTGACATTAAGATGGCCCTTAGGCTCAAAGATGGTGGGAGA<br>TACTTGGCCGACTTCAAGACCACTTACCGTGCTAAAAAGCCAGTGCAGATGCCAGGCGCATTCAATATCGACCGTAAGCTGGACATCA<br>CCTCGCATAATGAGGACTACACAGTCGTGGAACAATACGAGCGTTCTGTCGCTCGTCATTCTACCGGTGGATCTGGTGGTTCTATGGA<br>CTCTACAGAAGCGGTTATCAAAGAATTCATGCGATTCAAGGTATTGAATCGTTTATTGCTTCTGTGATGTTGTTGTTGCATTTGTGGCCTT<br>CGTTGAGTTTTATGATCTGATTTCGATTGATACAGTTGTTTTATGATTCTGTGTTGTTGTTGATGTTTCAATTGTTGCGTAATTGATTCGTT<br>TAGGAACTGAACTTTTACTGGTGATGATTTAGGTCCACATGGAAGGATCGATGAATGGCCACGAGTTCGAAATAGAAGGTGAAGGCGA<br>AGGTCGTCCATACGAGGGTACACAAACCGCAAAGCTTCGTGTGACCAAAGGTGGTCCATTGCCGTTTAGCTGGGACATCTTGTCTCCA<br>CAGTTTATGTATGGGTCCCGTGCCTTCACAAAGCACCCGGCCGATATTCCAGATTATTGGAAGCAGTCATTCCCTGAAGGCTTCAAATG<br>GGAACGTGTTATGAACTTCGAGGACGGCGGTGCCGTGTCAGTTGCACAAGATACAAGTTTGGAGGACGGCACATTGATCTACAAAGTC<br>AAGCTGAGAGGGACGAACTTCCCACCGGATGGGCCAGTCATGCAAAAGAAAACTATGGGCTGGGAAGCGTCTACCGAAAGGTAACAT<br>TCCTTAGTTACCTTTCTTTTCTTTTTCCATCATAAGTTTATAGATTGTACATGCTTTGAGATTTTTCTTTGCAAACAATCTCAGGTTGTACC<br>CTGAAGATGTGGTCCTCAAAGGTGACATCAAAATGGCGCTCCGTCTGAAAGACGGCGGAAGATATTTGGCGGATTTTAAGACGACCTA<br>TCGAGCCAAAAAGCCGGTCCAAATGCCTGGTGCCTTTAATATCGACAGAAAACTCGACATTACCTCTCATAACGAAAGTCTTC |
| gSAS 222Dsto | GAAGATGTACGTGAGAGATGGAGTGCTCACGGGAGATATCGAAATGGCTCTCTTTTGAAGACTTACGAAGATTATACCGTCGTCGAGC<br>AGTATGAGCGTTCAGTTGCTAGGCATAGCACTGGTGGTAGTGGTGGATCTTGAGCTTAAGTCTTCAAATGGAGGGTGGTGCTCATTATC<br>GTTGTGATTTTAGAACAACATAC |

| | |
|---|---|
| gSAS 223A | CTTACAACTGCTTTCCATTATGGAATTTGAAGACTTAATGGATTCTACCGAGGCCGTGATCAAAGAATTCATGAGGTTCAAGTCTTCAAA<br>TAGTACGTGATGTTTGAAGACTTAGGTAGCGGCGGCGGAGGTAGCGGTTCGAAGTCTTCAAATGATATCCTTACAACTGCTTTCCATTA<br>TGGAAATAGAGTTTTTGCTAA |
| gSAS 223Dnon | TGATATCCTTACAACTGCTTTCCATTATGGAAATAGAGTTTTTGCTAATTTGAAGACTTACGAAGATTATACCGTCGTCGAGCAGTATGAG<br>CGTTCAGTTGCTAGGCATAGCACTGGTGGTAGTGGTGGATCTGGAGGTAAGTCTTCAAAAAATACCCTGATAACATCCAAGATTATTTC<br>AAACAATCTTTTCCAAAGGG |
| gSAS 224Ba | GAAGACTTGTTCAAGGTGCACATGGAAGGCAGCATGAACGGACATGAGTTCGAGATCGAAGGTGAAGGTGAGGGTAGACCTTACGAG<br>GGAACTCAGACTGCTAAGCTCAGAGTGACAAAGGTGGACCGCTTCCGTTCTCTTGGGATATCCTTTCACCGCAGTTCATGTACGGCT<br>CTAGGGCTTTTACTAAGCACCCTGCTGATATCCCGGACTACTGGAAGCAATCTTTCCCAGAGGGATTCAAGTGGGAGCGTGTGATGAA<br>TTTCGAGGATGGTGGTGCTGTTTCTGTGGCTCAGGATACTTCTCTTGAGGACGGAACCCTCATCTACAAGGTTAAGCTCCGTGGAACTA<br>ACTTCCCTCCTGATGGACCTGTGATGCAGAAAAAGACCATGGGCTGGGAAGCTTCTACTGAGAGACTTTACCCTGAGGACGTTGTGCT<br>CAAGGGCGATATCAAGATGGCTCTCAGACTTAAGGACGGTGGACGTTACCTCGCTGATTTCAAGACTACCTACAGGGCCAAGAAACCT<br>GTGCAAATGCCTGGGGCTTTCAACATCGATAGGAAGCTCGATATCACCAGCCACAACGAGGATTACACTGTGGTTGAGCAGTACGAGA<br>GATCTGTGGCTAGACACTCTACTGGTGGAAGCGGAGGATCTATGGATAGCACTGAGGCTGTCATCAAAGAATTCATGCGTTTCAAAGTT<br>CATATGGAAGGTTCCATGAATGGGCACGAATTTGAAATTGAAGGCGAAGGCGAGGGACGTCCGTATGAAGGTACTCAAACAGCTAAGC<br>TGAGGGTTACCAAAGGCGGTCCTCTTCCTTTTTCCTGGGATATCTTGAGCCCTCAATTCATGTATGGCAGCAGGGCCTTCACCAAACAT<br>CCAGCAGATATCCCTGATTATTGGAAACAAAGCTTCCCGGAAGGGTTTAAGTGGGAGAGAGTCATGAACTTCAAGTCTTCAA |
| gSAS 224Bb | GAAGACTTCTTCGAAGATGGCGGAGCAGTGAGTGTTGCTCAAGACACTTCTTTGGAGGATGGGACTCTGATCTATAAGGTGAAGTTGC<br>GTGGGACCAACTTTCCTCCAGACGGTCCGGTTATGCAAAAGAAAACGATGGGATGGGAAGCCAGCACCGAAAGACTTTATCCAGAGGA<br>TGTGGTGTTGAAGGGTGACATTAAGATGGCCCTTAGGCTCAAAGATGGTGGGAGATACTTGGCCGACTTCAAGACCACTTACCGTGCT<br>AAAAAGCCAGTGCAGATGCCAGGCGCATTCAATATCGACCGTAAGCTGGACATCACCTCGCATAATGAGGACTACACAGTCGTGGAAC<br>AATACGAGCGTTCTGTCGCTCGTCATTCTACCGGTGGATCTGGTGGTTCTATGGACTCTACAGAAGCGGTTATCAAAGAATTCATGCGA<br>TTCAAGGTCCACATGGAAGGATCGATGAATGGCCACGAGTTCGAAATAGAAGGTGAAGGCGAAGGTCGTCCATACGAGGGTACACAAA<br>CCGCAAAGCTTCGTGTGACCAAAGGTGGTCCATTGCCGTTTAGCTGGGACATCTTGTCTCCACAGTTTATGTATGGGTCCCGTGCCTTC<br>ACAAAGCACCCGGCCGATATTCCAGATTATTGGAAGCAGTCATTCCCTGAAGGCTTCAAATGGGAACGTGTTATGAACTTCGAGGACG<br>GCGGTGCCGTGTCAGTTGCACAAGATACAAGTTTGGAGGACGGCACATTGATCTACAAAGTCAAGCTGAGAGGGACGAACTTCCCACC<br>GGATGGGCCAGTCATGCAAAAGAAAACTATGGGCTGGGAAGCGTCTACCGAAAGGTTGTACCCTGAAGATGTGGTCCTCAAAGGTGAC<br>ATCAAAATGGCGCTCCGTCTGAAAGACGGCGGAAGATATTTGGCGGATTTTAAGACGACCTATCGAGCCAAAAAGCCGGTCCAAATGC<br>CTGGTGCCTTTAATATCGACAGAAAACTCGACATTACCTCTCATAACGAAAGTCTTC |

| | |
|---|---|
| gSAS 226A | AAATACCCTGATAACATCCAAGATTATTTCAAACAATCTTTTCCAAAGGGTTTGAAGACTTTTCGATGGATTCTACCGAGGCCGTGATCA AAGAATTCATGAGGTTCAAGTCTTCAAAGAAGATGGAGGAATCTGTAACGCTCGTAATGATATCACTATGGAGGGAGATACATTTTACAA CAAAGTTCGTTTTTACGGAA |
| gSAS 232A | GAAGACTTAATGTCTGAACTTATTAAGGAGAATATGCATATGAAGCTTTACATGGAGGGTACTGTTGATAACCATCATTTCAAGTGTACA TCTGAAGGAGAGGGTAAACCTTACGAAGGTACTCAAACAATGAGAATCAAGGTTGTTGAGGGTGGACCTCTACCATTCGCTTTCGATAT CTTGGCTACATCTTTTCTTTATGGTTCTAAAACTTTTATCAACCATACACAGGGAATTCCCGATTTCTTCAAGCAATCTTTCCCTGAGGGT TTCACTTGGGAGAGAGTTACTACATACGAGGATGGTGGAGTTTTGACTGCTACTCAAGATACATCTCTTCAGGATGGATGTCTCATCTA CAATGTTAAGATAAGAGGAGTTAATTTCACATCTAACGGTCCTGTTATGCAAAAAAAGACTCTTGGTTGGGAGGCTTTCACAGAAACTCT TTACCCTGCTGATGGTGGTTTGGAGGGAAGAAATGACATGGCCCTTAAGCTGGTTGGTGGATCACATCTTATCGCAAATATCAAGACTA CTTACAGATCTAAGAAACCGGCTAAAAATTTAAAAATGCCAGGTGTTTATTACGTGGACTATCGGTTGGAGAGAATTAAGGAGGCTAACA ACGAGACTTATGTCGAACAACATGAAGTGGCTGTTGCTAGATATTGTGATTTGCCATCTAAACTTGGACACAAGCTCAATGGAGGTAAG TCTTC |
| gSAS 452 | GTGCATTGGTCTCAACATAGGTGTGAGCAAGGGCGAGGAGCTGTTCACCGGGGTGGTGCCCATCCTGGTCGAGCTGGACGGCGACG TAAACGGCCACAAGTTCAGCGTGTCCGGCGAGGGCGAGGGCGATGCCACCTACGGCAAGCTGACCCTGAAGTTCATCTGCACCACCG GCAAGCTGCCCGTGCCCTGGCCCACCCTCGTGACCACCTTCGGCTACGGCCTGATGTGCTTCGCCCGCTACCCCGACCACATGAAGC AGCACGACTTCTTCAAGTCCGCCATGCCCGAAGGCTACGTCCAGGAGCGCACCATCTTCTTCAAGGACGACGGCAACTACAAGACCC GCGCCGAGGTAACATTCCTTAGTTACCTTTCTTTTCTTTTTCCATCATCTATCAATTTCTTTGCGGAAATTTATTTGAAGCTGTAGAGTTAA AATTGAGTCTTTTAAACTTTTGTAGGTGAAGTTCGAGGGCGACACCCTGGTGAACCGCATCGAGCTGAAGGGCATCGACTTCAAGGAG GACGGCAACATCCTGGGGCACAAGCTTGAGTACAACTACAACAGCCACAACGTCTATATCATGGCCGACAAGCAGAAGAACGGCATCA AGGTAAGTTGTTACTTATGATTGTTTTCCTCTCTGCTACATGTATTTTGTTGTTCATTTCTGTAAGATATAAGAATTGAGTTTTCCTCTGAT GATATTATTAGGTGAACTTCAAGATCCGCCACAACATCGAGGACGGCAGCGTGCAGCTCGCCGACCACTACCAGCAGAACACCCCCAT CGGCGACGGCCCCGTGCTGCTGCCCGACAACCACTACCTGAGCTACCAGTCCGCCCTGTTCAAAGACCCCAACGAGAAGCGCGATCA CATGGTCCTGCTGGAGTTCCTGACCGCCGCCGGGATCCCTCCGTTGTTGAGACCAATGCAC |

| | |
|---|---|
| gSAS453 | GTGCATTGGTCTCAACATCCTCTGGTGTCAAAAGGCGAGGAGCTCTTCACGGGCGTCGTACCCATACTCGTTGAGCTTGATGGAGATG<br>TGAATGGACATAAGTTTAGTGTTAGCGGAGAGGGAGAAGGTAAATCCTGGTCCACACTTTTACGATAAAAACACAAGATTTTAAACTATG<br>AACTGATCAATAATCATTCCTAAAAGACCACACTTTTGTTTTGTTTCTAAAGTAATTTTTACTGTTATAGCAGGAGATGCAACCTATGGAA<br>AATTGACGTTGAAATTTATATGTACTACTGGTAAGCTGCCAGTTCCCTGGCCGACCCTCGTCACTACACTGTCCTGGGGGGTCCAATGC<br>TTCGCTAGATATCCTGATCATATGAAGCAACACGACTTTTTCAAAAGCGCCATGCCCGAAGGGTACGTTCAGGAACGAACGATTTTCTT<br>CAAAGATGACGGAAACTATAAGACGAGGGCGGAAGTAAAGTTTGAGGGAGACACCTTGGTAAATCGAATAGAATTGAAGGGAATCGAC<br>TTTAAGGAAGATGGGAATATACTAGGCCATAAGCTAGAGTATAATGCGATCCACGGGAACGTGTATATTACCGCGGACAAACAAAAAAA<br>TGGTATTAAAGCTAATTTTGGCCTCAACTGTAATATCGAGGATGGATCTGTTCAGTTAGCTGACCACTATCAACAAAATACACCGATAGG<br>TAAAGCAACTGTGTTTTAATCAATTTCTTGTCAGGATATATGGATTATAACTTAATTTTTGAGAAATCTGTAGTATTTGGCGTGAAATGAGT<br>TTGCTTTTTGGTTTCTCCCGTGTTATAGGTGACGGTCCCGTACTACTGCCCGACAATCACTACCTTAGCACCCAGTCTGCGCTATTCAAA<br>GATCCTAACGAGAAGCGGGACCATATGGTTCTTTTGGAGTTCTTGACCGCCGCGGGGATTACTTTAGGTATGGATGAGTTATACAAGTA<br>AGCTTTTGTTGAGACCAATGCAC |
| gSAS 454 | GTGCATTGGTCTCAACATAGGTGTGAGCAAGGGCGAGGAGCTGTTCACCGGGGTGGTGCCCATCCTGGTCGAGCTGGACGGCGACG<br>TAAACGGCCACAAGTTCAGCGTGTCCGGCGAGGGCGAGGGCGATGCCACCTACGGCAAGCTGACCCTGAAGTTCATCTGCACCACCG<br>GCAAGCTGCCCGTGCCCTGGCCCACCCTCGTGACCACCTTCGGCTACGGCCTGATGTGCTTCGCCCGCTACCCCGACCACATGAAGC<br>AGCACGACTTCTTCAAGTCCGCCATGCCCGAAGGCTACGTCCAGGAGCGCACCATCTTCTTCAAGGACGACGGCAACTACAAGACCC<br>GCGCCGAGGTGAAGTTCGAGGGCGACACCCTGGTGAACCGCATCGAGCTGAAGGGCATCGACTTCAAGGAGGACGGCAACATCCTG<br>GGGCACAAGCTTGAGTACAACTACAACAGCCACAACGTCTATATCATGGCCGACAAGCAGAAGAACGGCATCAAGGTGAACTTCAAGA<br>TCCGCCACAACATCGAGGACGGCAGCGTGCAGCTCGCCGACCACTACCAGCAGAACACCCCCATCGGCGACGGCCCCGTGCTGCTG<br>CCCGACAACCACTACCTGAGCTACCAGTCCGCCCTGTTCAAAGACCCCAACGAGAAGCGCGATCACATGGTCCTGCTGGAGTTCCTGA<br>CCGCCGCCGGGATCCCTCCGTTGTTGAGACCAATGCAC |
